# ER-phagy drives age-onset remodeling of endoplasmic reticulum structure-function and lifespan

**DOI:** 10.1101/2024.08.07.607085

**Authors:** EKF Donahue, NL Hepowit, B Keuchel, AG Mulligan, DJ Johnson, M Ellisman, R Arrojo e Drigo, J MacGurn, K Burkewitz

## Abstract

The endoplasmic reticulum (ER) comprises an array of structurally distinct subdomains, each with characteristic functions. While altered ER-associated processes are linked to age-onset pathogenesis, whether shifts in ER morphology underlie these functional changes is unclear. We report that ER remodeling is a conserved feature of the aging process in models ranging from yeast to *C. elegans* and mammals. Focusing on *C. elegans* as an exemplar of metazoan aging, we find that as animals age, ER mass declines in virtually all tissues and ER morphology shifts from rough sheets to tubular ER. The accompanying large-scale shifts in proteomic composition correspond to the ER turning from protein synthesis to lipid metabolism. To drive this substantial remodeling, ER-phagy is activated early in adulthood, promoting turnover of rough ER in response to rises in luminal protein-folding burden and reduced global protein synthesis. Surprisingly, ER remodeling is a pro-active and protective response during aging, as ER-phagy impairment limits lifespan in yeast and diverse lifespan-extending paradigms promote profound remodeling of ER morphology even in young animals. Altogether our results reveal ER-phagy and ER morphological dynamics as pronounced, underappreciated mechanisms of both normal aging and enhanced longevity.

## Introduction

The endoplasmic reticulum (ER) houses various cellular processes fundamental to cell homeostasis and healthy aging, including protein quality control, lipid and membrane biosynthesis, autophagy initiation, carbohydrate metabolism and calcium signaling^1,2^. The ER is also considered an integrator hub for coordinating intracellular signaling and the morphological dynamics of diverse other membrane-bound organelles through a complex and malleable array of inter-organelle contact sites^3,4^. However, the impact of the ER extends beyond the intracellular space. By integrating a variety of nutrient-sensing pathways with its functions in facilitating secretion, ER status is a key determinant of healthy intercellular signaling in metazoans^5,6^. This central positioning of the ER in cell and organismal physiology engenders vulnerability, where disruption of the ER can trigger widespread dysfunction from the subcellular to the organismal scale.

Indeed, dysregulation of virtually every ER-mediated process is associated with the chronic diseases of aging^7,8^. Understanding the mechanisms of age-onset ER dysfunction is thus critical for developing strategies to promote healthier aging.

Traditionally, the stress signaling and transcriptional pathways that comprise the ER unfolded protein response (UPR) have held the focus of the aging field^1,9^. Because the UPR activates expression of ER chaperones and genes capable of boosting protein-folding capacity and other ER functions, the response is generally considered protective. However, aging is associated with compromised ER function and dysregulated UPR signaling, including declining ER chaperone levels and reduced UPR inducibility^1,10–13^. Furthermore, chronically high baseline UPR signaling, as occurs during an unresolvable ER stress, is linked to enhanced inflammation and metabolic dysfunction commonly associated with advanced age^1,5,14,15^. Thus, aging appears to involve declining ER proteostasis functions in conjunction with aberrant ER stress signaling responses, which can elicit pleiotropic metabolic and immune consequences, but how this situation arises is not clear.

While these stress-signaling pathways are indeed critical during aging, ER morphodynamics represents an emergent, understudied mechanism for cells to shape ER metabolic outputs and maintain homeostasis. The essential functions of the ER are compartmentalized into distinct, structural subdomains^2,4,16,17^. ER sheets, often in parallel stacks, are the primary sites of protein synthesis and maturation. Sheets are typically heavily studded with ribosomes, giving the subdomain its signature rough appearance, and highly enriched with protein translocation and quality-control machinery, including signal recognition particle receptor and translocon subunits^2,4^. ER sheet morphologies are optimized for protein homeostasis by accommodating more polyribosome docking and supporting protein folding and maturation with larger luminal volume ratios^2,4^. On the other hand, highly curved ER tubular structures provide less ribosome docking and are more associated with lipid-droplet biogenesis and interactions with other organelles^2,8^. The relative abundance of sheet versus tubule subdomains varies widely between cell types and corresponds to the functional specialization of the cell. Extreme examples include pancreatic acinar cells, entirely filled with rough ER sheets needed for secreting massive amounts of digestive enzymes, and adipocytes, functionally dedicated to lipid metabolism and accordantly possessing predominantly tubular ER^2^. It is also increasingly clear that (patho)physiological fluctuations can trigger altered ER morphology within a given cell type^18–20^, and reciprocally, defects in ER-shaping factors are linked to the pathogenesis of neurodegeneration and metabolic disease^20–23^. However, whether ER morphological transitions are age-dependent and important in instigating age-related chronic diseases remains little explored.

Live cell and *in vitro* reconstitution assays have revealed the factors shaping ER subdomains to be dynamic and multifactorial. ER-shaping proteins are known to stabilize morphology via the relative abundances of sheet- and tubule-shaping proteins^4,24^ such as the conserved reticulon- and DP1/Yop1-family proteins, which utilize hairpin motifs within the membrane bilayer to stabilize highly curved tubule morphologies and sheet-edges^2,24,25^. Functional partners also affect ER shaping – for instance, extensive ribosome docking is critical for stabilizing ER sheet subdomains in at least some contexts^26^. Although less explored, an alternative pathway for ER remodeling involves the selective targeting of ER components or subdomains for degradation via ER-phagy. ER-phagy occurs in several ways, including direct endolysosomal engulfment of ER fragments via micro-ER-phagy and autophagosomal engulfment via macro-ER-phagy^27^. In yeast and tissue-culture models, studies have revealed diverse ER-associated receptors directing selective targeting of ER to autophagosomes^27^. Notably, past screens that led to the discovery of ER-phagy adaptors utilized nutrient deprivation and mTOR inhibitors^27–29^, interventions known for extending lifespan, as the contexts for inducing high-level ER-phagy. Furthermore, key ER-phagy pathways share some molecular machineries with macro-autophagy, which recurrently emerges as an essential process for diverse pathways of lifespan extension. While the importance of the autophagic machinery broadly is well established in aging, whether forms of selective autophagy or particular autophagic cargoes, such as ER, play critical roles remains underexplored in these contexts.

Here, through a combination of yeast, *C. elegans* and mammalian models, we reveal the morphological dynamics of the ER during normal aging and propose new roles for ER-phagy in lifespan determination. By visualizing ER subdomains across tissues *in vivo* with super-resolution and confocal imaging, we found that ER remodeling is among the earliest and most profound cell biological changes after animals reach adulthood. ER networks in diverse cell types of aged animals exhibit a substantial decline in ER mass, particularly of rough ER sheets, thus potentially revealing a cell biological explanation for the reduced proteostatic capacity widely observed during aging. ER remodeling is an active process driven by ER-phagy program that is activated in response to luminal protein burden and established declines in global protein synthesis with age. Finally, we show that ER-phagy mutants phenocopy the effects of macroautophagy impairment on chronological lifespan in yeast, and that diverse lifespan-extension paradigms in *C. elegans* pro-actively promote alternative ER morphologies in young animals, together indicating that ER-phagy is a protective mechanism. These findings illuminate important new roles for ER morphological dynamics in modulating age-dependent decline.

## Results

### In vivo imaging of ER dynamics in adult C. elegans

Within the contiguous ER network, many proteins are highly enriched in specific structural and functional subdomains^10,18,24^. To study *in vivo* ER dynamics during aging, we developed transgenic *C. elegans* strains with labels capable of distinguishing rough or tubular ER. SEC-61.B is an essential component of the ER translocon that is enriched in rough ER sheets^18^. Genomic, N-terminal *GFP* fusion produced a native marker, GFP::SEC-61.B, with enrichment in rough ER and relative exclusion from established smooth ER subdomains, such as muscle sarcoplasmic reticulum (SR) (Figure S1A). Importantly, using native markers circumvents the loss of subdomain fidelity associated with overexpressed and/or heterologous markers^18,25^ (Figure S1A). To visualize smooth and tubular ER subdomains we inserted mKate2 at the C-terminus of the sole *C. elegans* reticulon, RET-1. Reticulons are conserved hairpin-domain protein that stabilize membrane curvature in ER tubules and sheet edges^19^, and which are among the most highly enriched proteins in the smooth ER^10^. Super-resolution imaging of these endogenous GFP::SEC-61.B and RET-1::mKate2 proteins revealed canonical SEC-61.B-enriched rough ER sheets linked by a RET-1-enriched network of ER tubules (Figure S1B). To further validate the robustness of the labeling approach, we swapped fluorophores on RET-1 to GFP and employed an alternative translocon subunit^26^, TRAP-1::mCherry. Live, super-resolution imaging of *trap-1::mCherry; ret-1::GFP* animals again revealed distinct enrichment of RET-1 on sheet edges and tubules (Figure 1A). Overall, co-expression of tagged translocon subunits resulted in strong co-localization (Figure S1C-D) and each translocon-reticulon pair (TRAP-1::mCherry; RET-1::GFP and GFP::SEC-61.B; RET-1::mKate2) revealed consistent overall ER morphology, indicating localization is independent of fluorescence labels. Interestingly, the relative abundance of SEC-61.B or RET-1 labeling in different tissues also revealed the enrichment of rough versus smooth ER according to the functional specialization known for each cell type (Figure 1B). For example, neuronal projections, muscle SR, and smooth muscle-like spermathecae showed high enrichment of smooth ER tubules via high levels of RET-1, while hypodermis and intestine, which execute robust collagen and lipoprotein secretion, were highly enriched for SEC-61.B-labeled rough ER^24^ (Figure 1B). Together, these results indicate that native labeling of SEC-61.B and RET-1 provides a useful platform for monitoring *in vivo* ER subdomain dynamics across tissues in *C. elegans*.

**Fig. 1.**
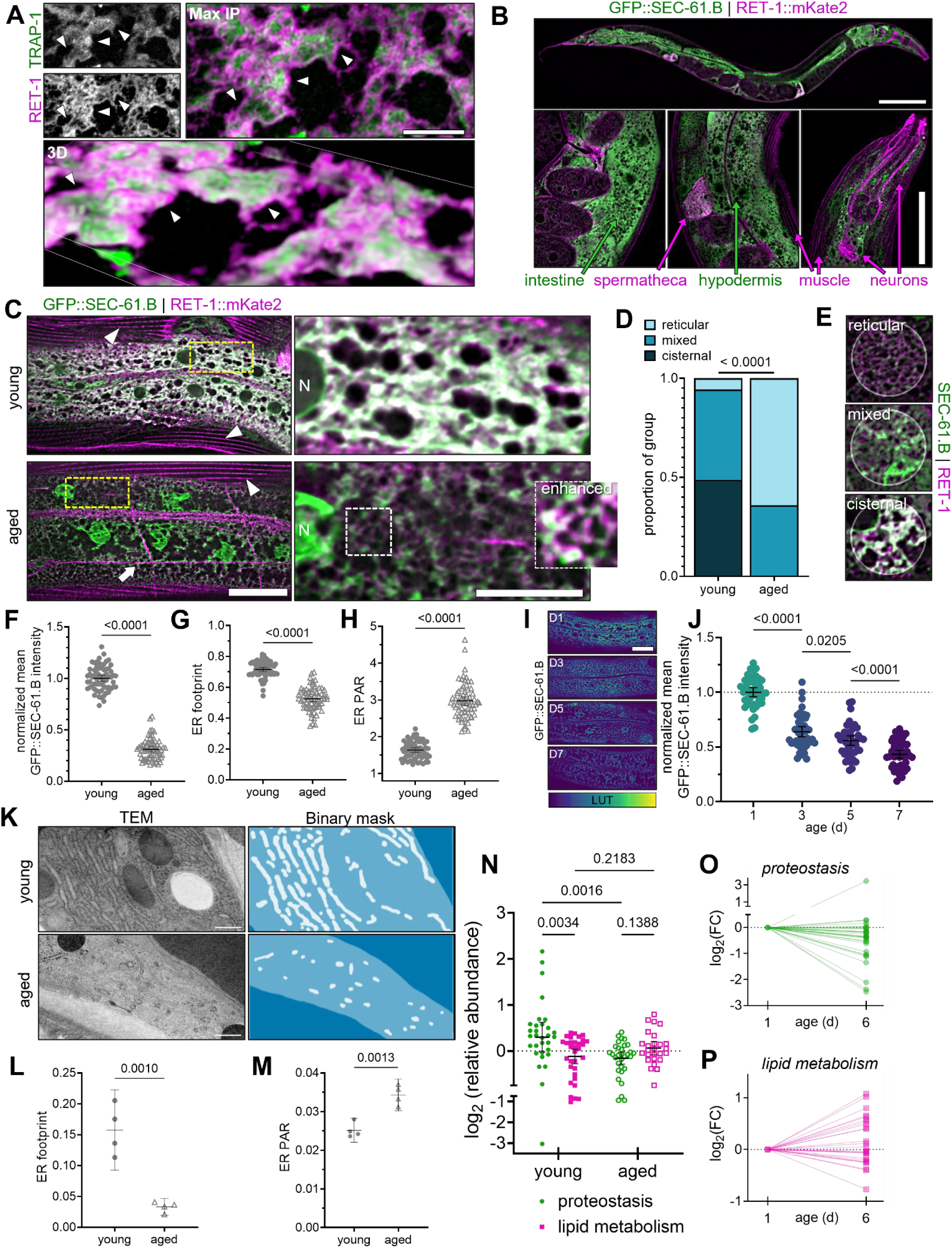
ER mass and morphology are highly dynamic during aging in *C. elegans*. (A) Live, super-resolution imaging of the ER marked by RET-1::GFP (ER tubules and sheet edges) and TRAP-1::mCherry (rough ER) in *C. elegans* hypodermis. Merged images represent maximum intensity projection (top right) and 3D projection (bottom). Arrows indicate RET-1::GFP enriched on the edges of TRAP-1-labeled rough ER sheets. Scale: 5 µm. (B) Fluorescence imaging of GFP::SEC-61.Β; RET-1::mKate2 depicts differential enrichment of ER subdomain markers between tissues. Top scale: 100 µm; bottom scale: 10 µm (C) Fluorescence imaging of GFP::SEC-61.Β; RET-1::mKate2 in young (day-1) and aged (day-7) adults. Arrows and arrowheads indicate smooth ER subdomains of neuronal projections and muscle SR, respectively. Zoom insets depict hypodermal ER network. Scale: 10 µm. (D, E) Categorical analysis of ER morphology (D) with reference images (E) for N = 123 regions from 42 young worms, 120 regions from 44 aged worms, pooled from 2 independent experiments. X^2^-test, p < 0.0001. (F-H) Quantification of normalized mean intensity (F), footprint (G), and perimeter:area ratio (H) in young vs. aged GFP::SEC-61.B worms. N = 64 young, 67 aged, pooled across 3 replicates. Analyzed via t-tests. (I, J) Representative images (I) and quantification (J) of a timecourse of GFP::SEC-61.B imaging in hypodermal cells. N = 45 worms/age, pooled across 3 replicates. One-way ANOVA, p < 0.0001, post hoc Šidák test. Scale: 10 µm. (K) TEM images (left) and binary masks (right, white) of the ER in hypodermis of young and aged worms. Scale: 500 nm. (L-M) Quantification of hypodermal ER footprint (L) and PAR (M). N = 4 worms/group. Analyzed via t-tests. (N-P) The ER proteome in young (day-1) and aged (day-6) worms reveals relative shifts in functional subdomain composition during aging. Data points represent relative abundance of a distinct peptide within each functional category (N_proteostasis_ = 31 young, 31 aged; N_lipidMetabolism_ = 32 young, 26 aged). Analyzed via mixed model, post hoc Fisher’s LSD test. Age-dependent trajectories displayed in O and P are normalized to abundance in young animals. Data derived from proteomic study by Walther et al. For all graphs, error bars indicate mean ± 95% CI. For images, “N” = nucleus.

### Aging is associated with declines in total ER mass and remodeling of ER structure-function

We next asked if ER morphology is dynamic over the worm’s lifespan. As a thin and metabolically flexible cell type, the *C. elegans* hypodermis is well suited for visualizing fine ER structures *in vivo* using GFP::SEC-61.B and RET-1::mKate2. In young day-1 adults, the ER appeared as a dense network of SEC-61.B-enriched cisternal structures extending throughout the cell (Figure 1C). In young animals, RET-1 was primarily associated with sheet-edges rather than resolvable RET-1-enriched ER tubules (Figure S1D, 1C). We then aged animals to day 7 of adulthood, a stage coinciding with the onset of age-dependent functional decline. Consistent with prior descriptions of both *C. elegans*^27^ and mammalian aging and progerias, the SEC-61.B-marked nuclear envelope developed significant structural distortion (Figure 1C)^28^. Focusing on the peripheral ER network, however, we found that these aged adults showed marked reduction of GFP::SEC-61.B-enriched sheets, instead forming small, sparsely distributed clusters connected by a RET-1-enriched tubular network (Figure 1C). Overall, the structure of the peripheral ER shifted with aging from a dense, cisternal morphology to a more diffuse and predominantly tubular network (Figure 1D-E).

We also tested this model more quantitatively and to ensure our observations were independent of specific fluorescence tags or co-expression of multiple tags within the ER. We imaged the ER in transgenic lines expressing GFP::SEC-61.B and RET-1::GFP alone (Figure 1F-H, Figure S1E-I). Aging revealed similar changes in ER morphology, as well as substantially reduced fluorescence signal from GFP::SEC-61.B and RET-1::GFP (Figures 1F, S1G), indicating a significant loss of both translocon and ER-shaping proteins with age. We confirmed these results independently of fluorescence imaging using epitope-tagged transgenic lines and immunoblotting (Figures S1J-M). To quantitate the decline in ER mass and shift from sheet to tubular morphologies, we analyzed footprint and perimeter:area ratios (PAR). We found that the ER footprint declined by ∼25% (Figures 1G, S1H), indicating that the overall organelle volume was reduced with aging, rather than simply possessing a reduced complement of SEC-61.B and RET-1 proteins. Finally, the ER perimeter:area ratio increased roughly two-fold for both reporters, consistent with a shift towards a more tubular network (Figures 1H, S1I). To determine the timeline of ER remodeling once animals reached adulthood, we imaged GFP::SEC-61.B animals across early-mid adulthood. The finding that fluorescence intensity decreased by 36% over only the first two days revealed that ER remodeling was a surprisingly early event in the aging process (Figure 1I-J). Finally, we employed transmission electron microscopy (TEM) to image the ER of unlabeled, wild-type animals. Consistent with our fluorescence imaging, young animals exhibited densely packed stacks of rough ER sheets that gave way to a sparse, tubular network (Figure 1K), further supported by reduced ER footprint and an increase in ER perimeter-area ratio (Figure 1L-M).

These age-dependent morphological shifts may correspond to functional shifts from proteostasis functions of the rough ER sheets to those that are more tubule-associated, such as lipid metabolism^2^. To support this model, we mined recent proteomic datasets examining the age-dependent proteome of *C. elegans*^29^. We identified ER-resident proteins associated with either proteostasis or lipid metabolism and compared their levels between young and aged animals (Figure 1N-P). Intriguingly, the proteostasis network of the ER underwent a pronounced decline, mirroring the loss of rough ER (Figure 1N,O). In contrast, the levels of ER-associated proteins involved in lipid metabolism generally stayed consistent and roughly half of these proteins increased over age (Figure 1N,P). Together, we conclude that ER morphological shifts mirror a functional shift from proteostasis to lipid metabolism, based on the pronounced declines in total ER abundance and shift in ER structure-function from tightly packed stacks of rough ER cisternae toward diffuse tubular networks.

### Age-dependent ER remodeling is conserved between tissues, sex, and species

While the anatomy and metabolic plasticity of the *C. elegans* hypodermis is optimal for *in vivo* imaging of ER dynamics, we also aimed to determine which age-dependent changes in the ER network might be generalizable across other cell types with distinct physiological functions. Consistent with the intestine’s many secretory functions, intestinal cells possess densely packed and predominantly rough ER networks^30^ highly enriched for SEC-61.B (Figure 2A). Similar to the hypodermis, the intestine experiences a dramatic loss of SEC-61.B at the protein level, a reduction in ER footprint, and an increase in perimeter-area ratio (Figures 2B-D). Despite very different physiological roles and subcellular architectures, muscle cells (Figures 2E-H) and neurons (Figures 2I-L) exhibit declines in SEC-61.B intensity, ER footprint, and shifts in ER morphology as measured in the muscle belly and soma, respectively. Overall, these results reveal that the age-dependent loss of ER mass and ER remodeling occurs across most major tissue types in *C. elegans*.

**Fig. 2.**
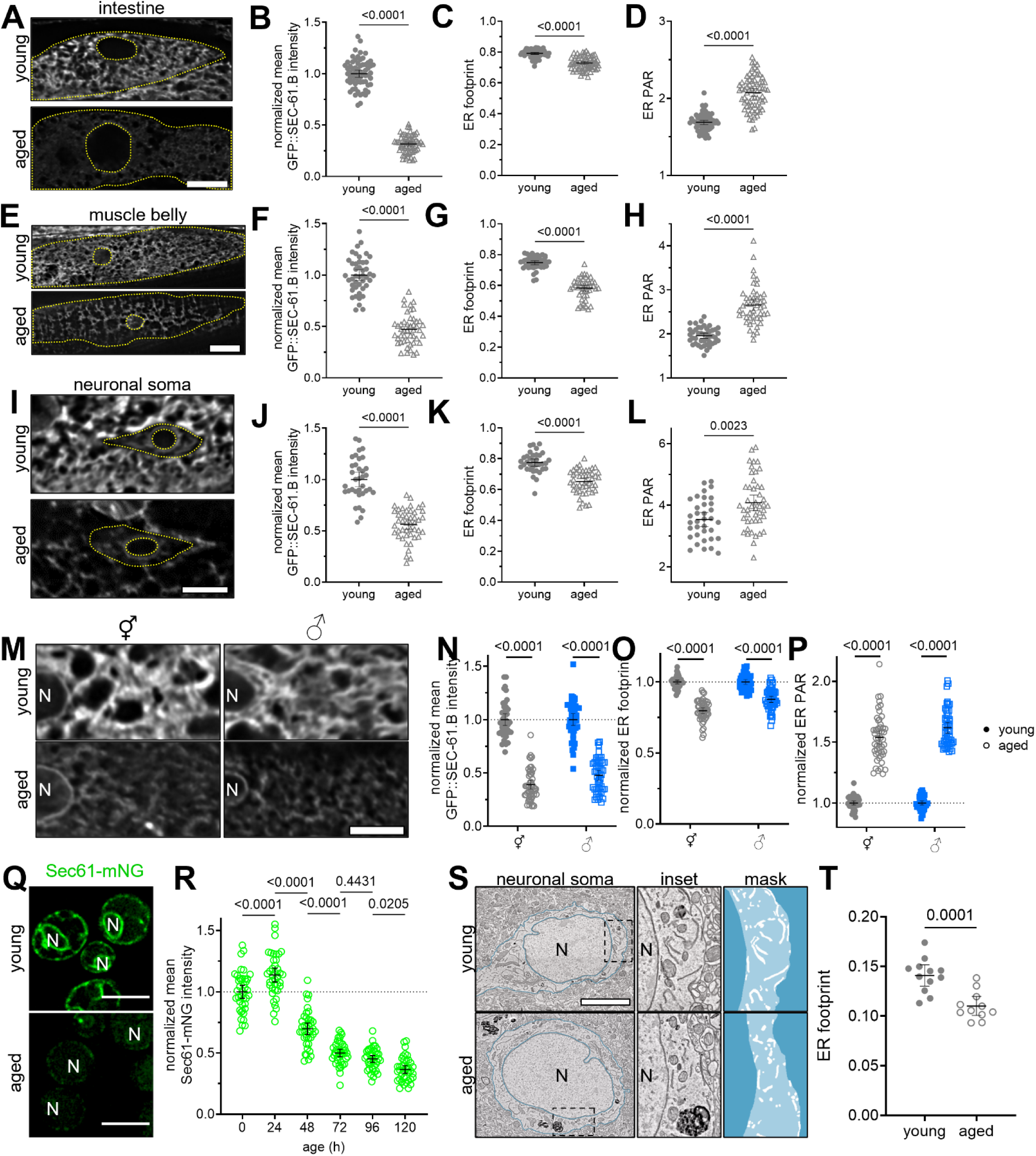
Age-onset ER remodeling occurs across tissues, sex, and species. (A) Imaging of intestinal GFP::SEC-61.Β in young and aged worms. Scale: 10 µm. (B-D) Quantification of normalized mean intensity (B), footprint (C), and PAR (D) of intestinal GFP::SEC-61.B in young and aged worms. N = 64 young, 70 aged, pooled across 3 replicates. Analyzed via t-tests. (E) Imaging of GFP::SEC-61.Β in the body wall muscle belly of young and aged worms. Scale: 10 µm. (F-H) Quantification of normalized mean intensity (F), footprint (G), and PAR (H) in muscle of young and aged GFP::SEC-61.B worms. N = 44 young, 47 aged, pooled across 3 replicates. Analyzed via t-tests. (I) Imaging of GFP::SEC-61.Β in the soma of the ALM neuron of young and aged worms. Scale: 10 µm. (J-L) Quantification of normalized mean intensity (J), footprint (K), and PAR (L) of GFP::SEC-61.B in the ALM soma of young and aged worms. N = 35 young, 46 aged, pooled across 3 replicates. Analyzed via t-tests. (M) Imaging of hypodermal GFP::SEC-61.Β in young and aged, hermaphrodite (⚥) and male (♂) worms. Scale: 10 µm. (N-P) Quantification of mean intensity (N), footprint (O), and PAR (P) of hypodermal GFP::SEC-61.B in young and aged worms, normalized to d1 for each sex. N_⚥_ = 41 young worms, 50 aged worms; N_♂_ = 43 young worms, 49 aged worms; pooled across 3 replicates. Analyzed via t-tests. (Q) Imaging of Sec61-mNG during chronological aging in young (0 h) and aged (72 h) yeast cells. Scale: 5 µm. (R) Sec61-mNG intensity during chronological aging in yeast. N = 40 cells/time point, pooled across 2 replicates. One-way ANOVA, p < 0.0001, post hoc Šidák tests. (S) SEM images of L2 motor cortex neurons in young (6 mo) and aged (18 mo) male mice with zoom insets (middle) and binary masks of ER (right, white). Scale: 2.5 µm. (T) Quantification of ER footprint in murine cortical motor neurons. N = 12 young cells, 11 aged cells. T-test, p = 0.0002. For all graphs, error bars indicate mean ± 95% CI. For images, “N” = nucleus.

While some aspects of age-dependent ER dynamics appear generalizable across many cell types, we also observed some changes to be highly tissue-specific. For example, RET-1::GFP strongly labels the young intestinal ER, but is virtually undetectable in aged intestine (Figures S2A and S2B), exhibiting a more pronounced decline than SEC-61.B in this tissue. Interestingly, given the morphological shift towards more tubular ER with age, the loss of RET-1 may suggest a shift towards alternative membrane-curving proteins, such as YOP-1. Additionally, muscle cells and neurons harbor specialized smooth ER subdomains within anatomically distinct regions, the myofilaments and neurites^30,36^, respectively. These smooth ER subdomains appear resistant to age-related ER declines, as the muscle SR exhibits only a ∼15% decline in RET-1::GFP intensity and ∼8% decline in SR footprint (Figures S2C-E) compared to ∼55% and ∼22% declines in the ER of the muscle belly (Figures 2E-H). Contrasting with every other tissue, RET-1 levels in neuronal projections generally appeared to be elevated. Specifically, we measured ∼two-fold increases in RET-1::GFP intensity with age in representative sensory (ALN/PLM) and motor (DA7) neurons (Figure S2G), while cytosolic GFP levels remain static, indicating that the age-related increases in RET-1 are independent of global neuron remodeling (Figure S2H). Collectively, the shift towards more tubular morphologies observed across cell types and the divergent dynamics of these specialized smooth ER subtypes suggest a model where rough ER subdomains are preferentially targeted for turnover as animals age.

Finally, to determine whether age-onset remodeling of ER is sex-dependent and evolutionarily conserved, we examined ER network changes in male *C. elegans*, yeast and mammalian systems. First, we found that males exhibit similar age-dependent declines in SEC-61.B intensity, footprint, and morphology in both the hypodermis (Figures 2M-P) and intestine (Figures S2L-O), revealing that the decline in ER is independent of egg production and the excessive venting of yolk associated with hermaphrodite aging^37^. Similar to *C. elegans,* we found that yeast cells experience progressive declines in Sec61-mNG beginning early during chronological aging (Figures 2Q and 2R). Lastly, scanning EM analysis of the peripheral ER network of 18-month mouse cortical neurons also revealed significant declines in ER mass (Figures 2S and 2T). Altogether, these results indicate that age-dependent ER remodeling is a conserved phenomenon.

### ER-phagy drives age-onset ER remodeling

Next, we set out to identify mechanisms promoting ER remodeling during aging. Given pronounced loss of both ER protein and membrane mass (Figure 1, S1), we reasoned that autophagic and/or lysosomal degradation processes are involved. To test whether autophagy drives ER loss, we fed animals dsRNA targeting both the upstream initiator ULK-1/Atg1/*unc-51* and downstream cargo adaptor GABARAP/Atg8/*lgg-1.* Autophagy inhibition via genetic depletion of Atg1/*unc-51* and Atg8/*lgg-1* had few discernible effects on ER size and shape in young adult animals, indicating that autophagosomal turnover plays a limited role in shaping the ER during development. However, knockdown of either Atg1/*unc-51* or Atg8/*lgg-1* suppressed virtually all age-onset changes. In particular, Atg8/*lgg-1(RNAi)* completely abrogated the loss of ER footprint and intensity of both GFP::SEC-61.B (Figures 3A-D) and RET-1::GFP (Figures 3E-H). Inhibiting autophagy via Atg8/*lgg-1(RNAi)* also preserved GFP::SEC-61.B and RET-1::GFP levels in the intestine, suggesting autolysosomal ER turnover drives ER remodeling across multiple tissues (Figures S3A-H).

**Fig. 3.**
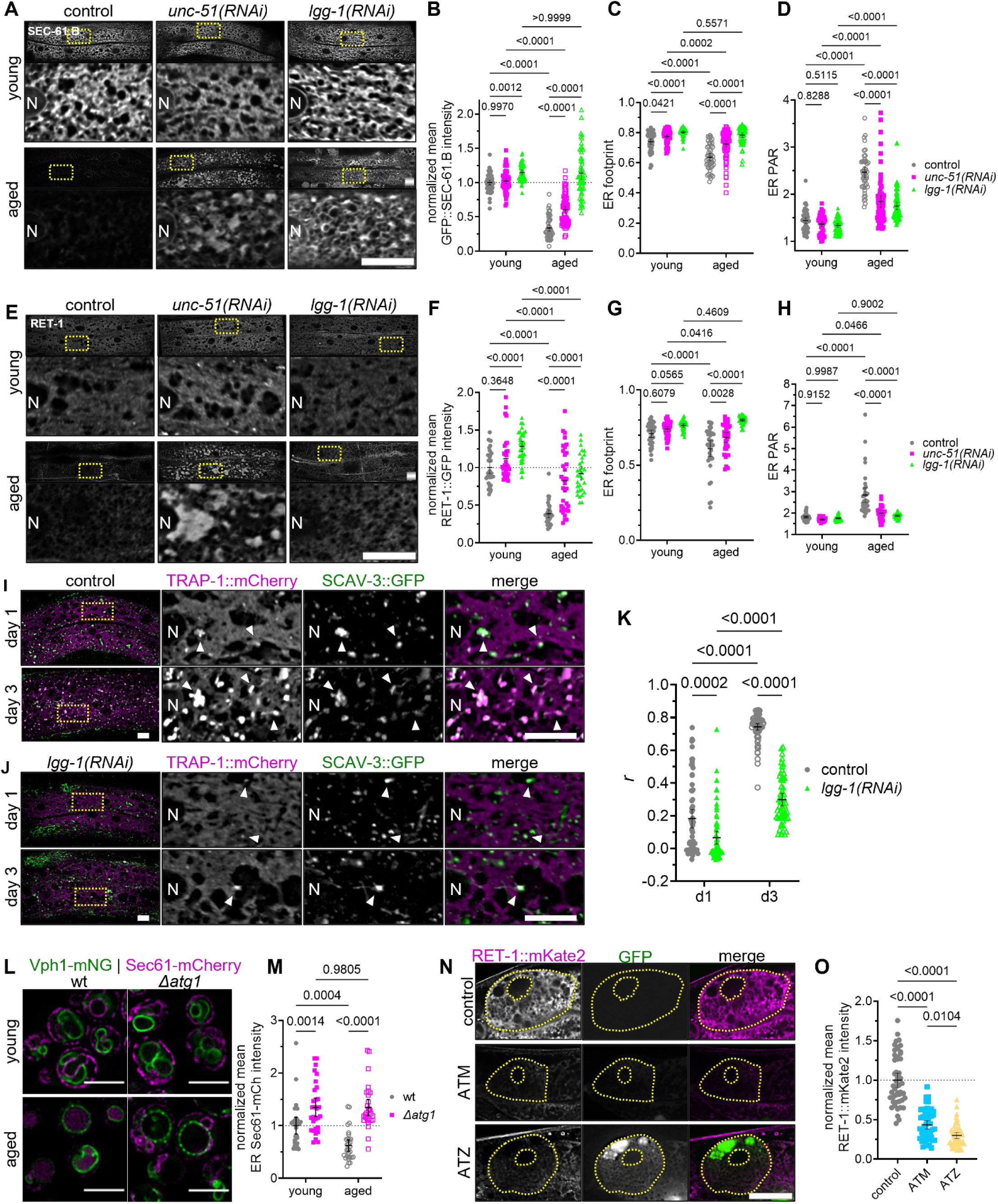
ER-phagy drives age-dependent ER remodeling. (A) Imaging of hypodermal GFP::SEC-61.Β in young and aged control, *unc-51(RNAi)*, and *lgg-1(RNAi)* worms. Scale bars: 10 µm. (B-D) Hypodermal GFP::SEC-61.B normalized mean intensity (B), footprint (C), and PAR (D) in young and aged control, *unc-51(RNAi)*, and *lgg-1(RNAi)* worms. Young = filled shapes; aged = hollow shapes. N_e.v._ = 57 young, 60 aged; N*_unc-51_* = 60 young, 67 aged; N*_lgg-1_* = 54 young, 56 aged; pooled across 3 replicates. Analyzed via two-way ANOVAs and post hoc Šidák tests. (E) Hypodermal RET-1::GFP in in young and aged control, *unc-51(RNAi)*, and *lgg-1(RNAi)* worms. Scale bars: 10 µm. (F-H) Hypodermal RET-1::GFP normalized mean intensity (F), footprint (G), and PAR (H) in young and aged control, *unc-51(RNAi)*, and *lgg-1(RNAi)* worms. Young = filled shapes; aged = hollow shapes. N_e.v._ = 35 young, 35 aged; N*_unc-51_* = 35 young, 35 aged; N*_lgg-1_* = 35 young, 31 aged; pooled across 2 replicates. Analyzed via two-way ANOVA and post hoc Šidák tests. (I-J) Hypodermal TRAP-1::mCherry (magenta) and SCAV-3::GFP (green) in d1 and d3 worms fed control (e.v., I) and *lgg-1* RNAi (J). Arrowheads mark regions of colocalization. Scale bars: 10 µm. (K) Pearson’s correlation coefficient (*r*) between mCherry and GFP in the hypodermis of control and *lgg-1(RNAi)* worms at d1 and d3. N_e.v._ = 65 d1, 69 d3; N*_lgg-1_* = 63 d1, 58 d3; pooled across 3 replicates. Two-way ANOVA, p < 0.0001, post hoc Tukey test. (L) Imaging of Sec61-mCherry and Vph1-mNG in young and aged control and *Δatg1* mutant yeast cells. Scale: 5 µm. (M) Mean cytoplasmic Sec61-mCherry intensity in control and *Δatg1* yeast at 0 h and 72 h. N = 30 yeast/group, pooled across 2 replicates. Two-way ANOVA, p < 0.0001, post hoc Tukey test. (N) Fluorescence imaging of intestinal RET-1::mKate2 in d1 adult control, GFP::ATM-, and GFP::ATZ-expressing worms. Dotted lines indicate cell membrane and nuclear envelope. Scale: 10 µm. (O) Quantification of intestinal RET-1::mKate2 intensity in control, GFP::ATM-, and GFP::ATZ-expressing worms, relative to controls. N = 51 worms/group, pooled across 3 replicates. One-way ANOVA, p < 0.0001, post hoc Tukey test. For all graphs, error bars indicate mean ± 95% CI. For images, “N” = nucleus.

The ER is an important source of membrane for autophagosome formation as well as a potential target of autophagosomes via ER-phagy, and both of these roles could potentially result in reduced ER mass^27,38,39^. To determine whether the ER is itself targeted as cargo for ER-phagy, we employed TRAP-1::mCherry. Unlike GFP and mKate2 labels, mCherry is resistant to lysosomal acidity and degradation, enabling analysis of lysosomal targeting^40^. In young adult animals, TRAP-1::mCherry and SEC-61.B::GFP labels reveal the same ER network organization (Figure S1B). However, TRAP-1::mCherry uniquely begins accumulating in distinct puncta by day 3 of adulthood (Figure 3I). Consistent with the age-onset formation of these puncta, we observed low colocalization of TRAP-1::mCherry with lysosomal membrane marker, SCAV-3::GFP, in animals on the first day of adulthood, and a dramatic increase in internalization of TRAP-1::mCherry within SCAV-3^+^ lysosomes by day 3 (Figures 3I-K, S3I). Depletion of Atg8/*lgg-1* prevented the formation of TRAP-1 puncta and suppressed lysosomal targeting (Figures 3J-K). Overall, these results are consistent with a model where activation of ER-phagy early in adulthood promotes turnover and remodeling of the ER. The role of autophagy in driving age-onset ER loss is evolutionarily conserved, as the decline of Sec61 in yeast correlates with re-localization to the vacuole via an Atg1-dependent mechanism (Figures 3L-M).

We hypothesized at least two major and non-exclusive driving forces for the age-onset targeting of ER for turnover. First, protein translation and ribosome abundance decrease during aging^35,41^, and thus reduced ribosome docking and protein flux through the ER could result in ER-phagy-mediated culling of rough ER subdomains. Secondly, global defects in the protein folding network are well known during aging, and if unfolded and/or aggregated proteins accumulate in the ER lumen, this would also promote ER-phagy^42–46^. To test these possibilities, we employed a model of enhanced ER secretory burden based on α1-antitrypsin fused with GFP^47^. Heterologous overexpression of wild type (ATM) or misfolding-prone α1-antitrypsin variant (ATZ) both add an exogenous secretory load, while ATZ also forms large intraluminal aggregates^42,47^ (Figure 3N; Figure S3J). We predicted that if reduced ER protein translocation drives age-onset loss of ER, then heterologously driving enhanced secretion of healthy ATM might delay or rescue declines in ER mass. Conversely, if luminal protein damage drives age-onset ER turnover, then elevated secretory burden (i.e., accumulating ATZ) would accelerate ER loss. Intriguingly, we found that overexpression of both ATM and pathogenic ATZ results in early reductions of ER mass (Figure 3O). These results support a model where age-associated ER turnover acts as an adaptive response to rising luminal protein folding burden, though age-dependent decreases in ribosome density and protein synthesis rates cannot be ruled out.

### ER remodeling is a common feature of lifespan extension

While the macroautophagy machinery is essential for longevity assurance in virtually all contexts tested, much less is known about whether forms of selective autophagy are important in lifespan determination. To determine whether ER-phagy indeed plays a role in adapting to age-onset stressors and promoting healthy lifespan, we first turned to yeast, where previously identified ER-phagy-specific adaptors allow us to uncouple the roles of ER-phagy and macroautophagy. We compared the chronological lifespans of wild type yeast with macroautophagy-deficient *Δatg5* mutants and ER-phagy-deficient *Δatg39*, *Δatg40*, and *Δsec66* mutants^48^. We found in each case that the ER-phagy mutants closely phenocopy the macroautophagy-deficient *Δatg5* mutant by reducing lifespan (Figure 4A-B, Table ST1), suggesting that the ER represents a critical but little explored target of the autophagic machinery in aging contexts.

**Fig. 4.**
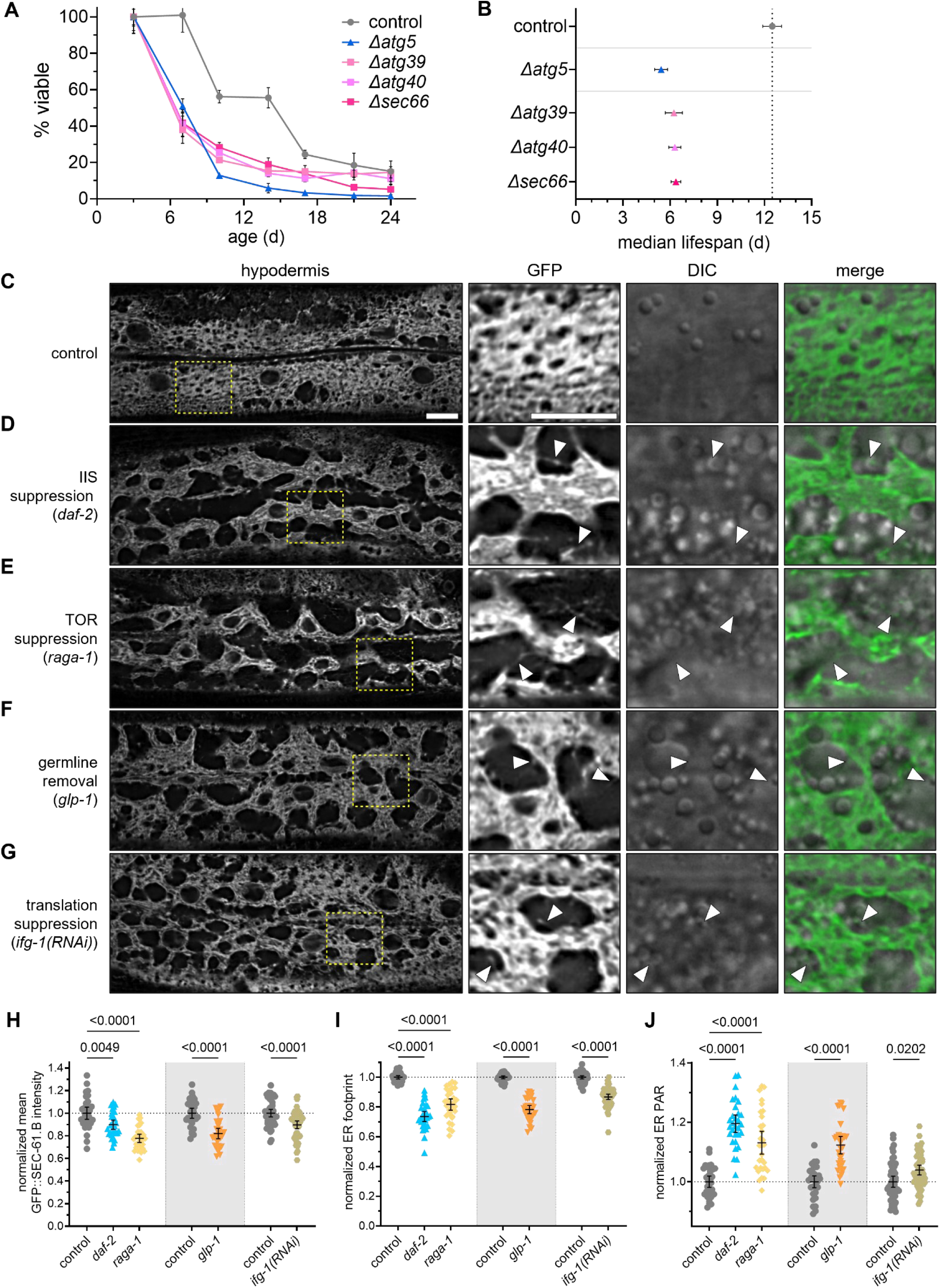
ER remodeling is a common feature of diverse lifespan extension paradigms. (A-B) Autophagy and ER-phagy are required for yeast chronological lifespan. (A) Percent of viable wild type and autophagy (*Δatg5* (blue squares)) or ER-phagy (*Δatg39*, *Δatg40*, *Δsec66* (pink triangles)) mutant yeast, normalized to d3. N = 4 biological replicates / strain. (B): median lifespan for each strain. One way ANOVA, p < 0.0001, post hoc Šidák test. See supplemental table ST1 for lifespan statistics. (C-G) Confocal imaging of hypodermal GFP::SEC-61.Β in young worms from control (N2, C), insulin/insulin-like signaling (IIS) suppression (*daf-2*, D), TOR suppression (*raga-1*, E), germline removal (*glp-1*, F), and translation suppression *(ifg-1(RNAi)*, G) conditions. DIC = differential interference contrast. All scales: 10 µm. Arrows indicate sparse ER tubules. (H-J) Quantification of mean intensity (H), footprint (I), and PAR (J) of GFP::SEC-61.B relative to wild type/controls. N = 30 control, 30 *daf-2*, 30 *raga-1*; 30 control_(25°C)_, 30 *glp-1*; 50 control_(e.v.)_, 52 *ifg-1(RNAi)*; pooled across 3 replicates. Colored backgrounds group experimental conditions with their respective controls. One way ANOVA, p < 0.0001, post hoc Šidák test. For all graphs, error bars indicate mean ± 95% CI.

We next sought to understand whether established paradigms of lifespan extension involve ER remodeling. We selected a panel of mechanistically diverse interventions, including reduced insulin/insulin-like signaling (IIS; *daf-2(e1370)*), inhibition of mTOR (*raga-1(ok386)*), germline removal (*glp-1(e2141)*), and inhibition of translation (*ifg-1(RNAi)*). Consistent with a model where ER remodeling is part of an adaptive process to age-onset stressors, we found that these interventions universally induce dramatic ER remodeling even at the onset of adulthood (Figures 4C-G). Long-lived worms exhibit substantial reductions in GFP::SEC-61.B intensity and footprint as well as increased ER perimeter-area ratio (Figures 4H-J). While some gross morphological changes resemble normal aging, we observe a number of unique forms of ER remodeling in long-lived animals. For example, in contrast to the evenly distributed rough ER sheets under normal conditions, the cisternal ER networks are strongly condensed into perinuclear regions, and large peripheral regions devoid of ER sheets are populated by sparse tubular networks (Figures 4B-F). This organization suggests enhanced functional compartmentalization of ER networks into distinct sheet-vs. tubule-filled regions of the cell, highlighting a need to visualize subcellular architecture in these contexts more globally. Feeding animals Atg8/*lgg-1* dsRNA to impair autophagy partly or fully restores GFP::SEC-61.B intensity, footprint, and PAR back to levels comparable to controls (Figures S4A-D), indicating that ER remodeling in long-lived animals also involves autophagy. However, the regionalized clustering of ER sheets persists during autophagy impairment (Figure S4A), indicating that this aspect of ER network remodeling in long-lived animals is independent of autophagy. Overall, these findings reveal that diverse interventions that promote lifespan extension induce substantial remodeling of the ER, likely in conjunction with associated organelle networks.

## Discussion

The morphological dynamics of the ER have received little attention compared to other organelles in aging contexts. Here, we established new tools in *C. elegans* for high-resolution, live-imaging of ER networks in aging metazoans, which revealed profound shifts in ER network morphology that are driven by ER-phagy. From surveying a variety of somatic tissues, we consistently found a decrease in ER protein levels, cellular ER volume, and a structural shift from densely packed sheet morphologies to diffuse and tubular networks (Figures 1, 2). A similar ER-network remodeling also occurs in yeast and mammalian neurons (Figures 2Q-T), highlighting an evolutionarily conserved aspect of the aging process relevant to diverse cell types. Finally, we found that ER-phagy drives turnover and remodeling of the ER network in both yeast and *C. elegans* while also playing an important role in yeast lifespan, thus newly linking this form of selective autophagy to aging biology. Identification of ER-phagy-specific receptors in *C. elegans* and other metazoan models would allow experimental uncoupling of ER-phagy from macroautophagy and further enable mechanistic dissection of the relative contributions of nonspecific macroautophagy and ER-phagy in diverse aging contexts.

As with any new age-associated phenomenon, primary concerns include determining whether the observed alteration is adaptive vs. maladaptive and whether it is causal or correlative to healthy aging. We propose a model where age-dependent ER remodeling serves as a causal, adaptive process associated with reprogramming of the proteostasis network^35,49^. This model is based on our findings that i) luminal protein damage triggers early loss of ER; ii) the ER proteome shifts away from proteostasis machineries with age; iii) ER-phagy-dependent turnover of ER promotes yeast chronological lifespan; and iv) long-lived animals adopt distinct, autophagy-dependent ER network configurations. While the net effect of ER-phagy on lifespan in yeast is positive (Fig. 4), we speculate that such early, pronounced remodeling of ER structures is likely to trigger a broader cascade of pleiotropic side-effects later in the aging process, especially in longer-lived cells and animals. For instance, a number of late-stage hallmarks of aging are known in other contexts to be strongly influenced by ER contributions. Examples of these hallmarks include declines in autolysosomal functions^7^, where the ER is a key source of membrane for both autophagosome biogenesis^50^ and lysosome repair^51^, and mitochondrial dysfunction^7^, where ER tubules are an emergent scaffold for regulating mitochondrial fission-fusion processes and biogenesis^3^. Further consistent with this model of ER remodeling as a potential trigger for additional age-associated changes in the cell, substantial ER remodeling becomes apparent prior or coincident with virtually all established hallmarks of aging in *C. elegans.* The early remodeling of the ER thus suggests a new example of antagonistic pleiotropy at the cell biological scale, where early-life ER transitions that may support remodeling of the cellular proteostasis network can trigger pleiotropic, off-target consequences later in life, after evolutionary pressures to enhance cell and organismal fitness have faded^52^. More broadly, given the delicate regulation of inter-organelle interactions needed to maintain metabolic fitness and the ER’s role as the hub of these interactions, our results suggest that changes in the subcellular architecture may represent an underappreciated form of age-related “damage”^53^. Similarly, the finding that diverse longevity paradigms trigger remodeling of the ER (Fig. 4) in conjunction with multiple other organelle networks^54,55^ suggests that cells may reconfigure the organelle ‘interactome’ in ways that shape the aging process.

Our results align well with several recent findings, while providing a new dimension for future studies. First, prior studies reported loss of the steady-state activity and inducibility of XBP-1 in *C. elegans* at the same early ages that we measured a substantial loss of ER mass^10,56^. The mechanisms linking these correlated changes remain unclear, but constitutive XBP-1 activation extends lifespan^57^ partly through a mechanism involving activation of lipophagy^43,58^, suggesting that XBP-1 could play an important role in coordinating these inter-related shifts in proteostasis, lipid metabolism and ER structure. The link between ER remodeling and the balance of protein:lipid metabolism is also apparent in mammals. High-fat diets in mice cause shifts from sheet-enriched to more tubular ER networks in the liver, and preventing these ER alterations by genetically enforcing sheet formation ameliorates obesity-related metabolic dysregulation^20^. Furthermore, aging is commonly associated with a rise in ectopic lipid accumulation in diverse cell types^59,60^, which our results suggest may similarly stem from shifted ER structure-function. The number of molecular and pathophysiological commonalities between obesity and aging, both producing increased risk of diverse age-onset disease^61^, suggest the similar sheet-to-tubule transitions in aged *C. elegans* and obese rodents could be a common etiological link. The genetic connections between ER-shaping factors and hereditary spastic paraplegia further support the potential sufficiency of altered ER morphology in driving disease in the nervous system as well^21,22^. Thus overall we provide foundational evidence supporting ER structure-function as an important new facet of geroscience, highlighting the potential for ER dynamics to serve as a targetable aging process underlying diverse forms of age-dependent pathophysiology.

## Acknowledgments

We thank the Burkewitz lab members for constructive feedback and B. Jacquet-Cribe, E. Diao, H.L. Singkhek, A.J. Kyle, and A. Pefanco for technical assistance. We thank A. Arruda and G. Parlakgul for technical advice, C. Wright for constructive feedback, and V. Gama, M. Patel, W. Mair and D. Miller for sharing reagents and equipment. The Caenorhabditis Genetics Center (NIH Office of Research Infrastructure Programs P40 OD010440) provided worm strains used in this work in addition to BQ5 from P. Hu, VK1882 and VK1950, provided by S. Pak and G. Silverman, and XW8056 provided by X. Wang. TEM sample preparation was performed through the Washington University Center for Cellular Imaging (WUCCI) at the Washington University School of Medicine and TEM imaging through the Vanderbilt Cell Imaging Shared Resource (supported by NIH grants CA68485, DK20593, DK58404, DK59637 and EY08126). This work was supported by the Glenn Foundation for Medical Research/American Federation for Aging Research, NIH/NIA R00AG052666 and R01AG073354 (KB), F31AG076290 and NIGMS T32GM007347 (ED), and R35GM144112 (JM).

## Author contributions

Conceptualization: ED and KB. Methodology: ED, NH, JM, and KB. Investigation: ED, NH, BK, AM, DJ, RD, and KB. Writing – Original Draft: ED and KB. Writing – Review & Editing: all authors. Funding Acquisition: ED and KB. Resources: ME and RD. Supervision: JM and KB

## Data and resource sharing

Worm strains generated in this study will be available on request and/or deposited to the Caenorhabditis Genetics Center (CGC) repository as appropriate. Yeast strains generated in this study will be available on request. This paper does not report original code. Any additional information required to reanalyze the data reported in this paper is available upon request. Further information and requests for data, resources, and reagents should be directed to and will be fulfilled by the lead contact, Kristopher Burkewitz.

## Supplemental information

Document S1. Figures S1–S4, tables TS1-3, and supplemental references.

## Methods

### C. elegans *husbandry*

Worms were grown and maintained on 6 cm nematode growth media (NGM)^62^ plates seeded with *E. coli* (OP50-1) at 20 °C*. E. coli* were cultured from single colonies in LB media overnight at 37 °C, after which 100 µL liquid culture were seeded per NGM plate to grow for two days at room temperature prior to use. For all experiments, worms were synchronized via timed egg-lay. In experiments involving *glp-1* mutants, 24 hours after synchronization *glp-1* mutants and controls were shifted to 25 °C for 24 h to impair germline development. See Supplemental Table 2 (ST2) for a complete list of worm strains used in this study.

### S. cerevisiae husbandry and chronological lifespan experiments

Yeast cultures in synthetic complete dextrose (SCD) medium were grown to mid-log phase (young cells) or chronologically aged to 72 hours from mid-log phase (aged cells) at 30 °C with agitation (220 rpm). See Supplemental Table 2 (ST2) for a complete list of yeast strains used in this study.

### M. muris husbandry and approval

Male FVB/NJ mice were obtained from the Jackson Laboratory (Bar Harbor, ME) and then were housed in cages of five mice and maintained on a 12-hour light/dark cycle with controlled temperature (21°C) and humidity. The mice had ad-libitum access to chow food (Rodent Diet 20 5053, PicoLabâ) and water. All animal procedures were approved by the Institutional Animal Care and Use Committee (IACUC) of Vanderbilt University (M2000086-00/01).

### RNAi experimental details

Gene-specific RNAi feeding clones (HT115) were obtained from the Ahringer RNAi Library (Source Bioscience), sequence-verified, and grown as described above for OP50-1 except the addition of 100 µg/mL carbenicillin (final concentration) to the LB and NGM. At least an hour prior to use, control and experimental lawns were spotted with 100 µL IPTG solution (0.1 M IPTG, 100 µg/mL carbenicillin, 12.5 µg/mL tetracycline) to induce dsRNA expression. For *ifg-1* RNAi, feeding began at the L4 developmental stage^63^. For *lgg-1* and *unc-51* RNAi aging experiments, the parental generation was also fed dsRNA from hatch for maximum penetrance. Otherwise, experimental worms were fed dsRNA from hatch.

### In vivo confocal microscopy – C. elegans

Synchronized worms were raised as described above. For each condition, an agarose mounting pad (10% agarose in M9 buffer^62^ was prepared prior to imaging by briefly heating the agarose solution and flattening it between two slides (#1.5, VWR). Approximately 3 µL 0.1 µm Polybead microsphere suspension (Polysciences) was added to the pad, worms were manually picked into the bead solution, and a coverslip (#1.5, VWR) was added to immobilize the worms. Imaging was performed on an Eclipse Ti2 inverted microscope with a Yokogawa CSU-W1 spinning disc (Nikon), a Plan Apo λ objective (10×/0.45 or 100×/1.45) (Nikon), and a Prime 95B sCMOS camera (Teledyne Photometrics). Type B and LDF immersion oils (Cargille Labs) were used, with the same type being used for all replicates within an experiment. Fluorophores were excited at 488 and 561 nm (dichroic mirror Di01-T405/488/568/647-13×15×0.5 (Semrock)), with respective ET525/36m and ET605/52m emission filters (Chroma). Additionally, DIC images (89101m emission filter) were captured for each fluorescent image. For all tissues, focal planes were selected at nuclear midlines. The ER was imaged in the same cells of each tissue across all conditions to account for potential cell-cell variability in ER networks, specifically anterior region of the hypodermis (*hyp7*), the first and last intestinal rings (*int1* and *int9*), body wall muscles, and the ALN neuron. A plane was selected within the myofilament lattice to image the SR. For neural ER projections, imaging was acquired via Z-stack, with analysis performed on maximum intensity-projections of the ALM, DA7, and PLM.

### In vivo microscopy – S. cerevisiae

Prior to imaging, yeast were concentrated via centrifugation at 3500 × *g*, pipetted onto a slide, and immobilized with a cover slip. Images were acquired with a DeltaVision Elite Imaging system [Olympus IX-71 inverted microscope; Olympus 100× oil objective (1.4 NA); DV Elite sCMOS camera, GE Healthcare]. Images were obtained on red (Alexa Fluor 594; 475 nm excitation, 523 nm emission), green (FITC; 575 nm excitation, 632 nm emission), and DIC channels.

### Image processing and data analysis

NIS-Elements Advanced Research (NIS-AR, Nikon) was used to process images for all worm experiments. Prior to analysis, images were subjected to automated 2-D or 3-D deconvolution and subsequent denoising via the Denoise.ai module. Rolling-ball background removal was performed on each color channel with identical settings for all images in the experimental replicate. For categorical analysis of ER morphology, images were blinded, and the JavaScript command “Math.random();” was used within NIS-AR’s GA3 module to select 3 random regions within the hypodermal ROI. Subregion morphology was categorized using the Qualitative Annotations plugin in Fiji^64^. For ER intensity and morphological quantitation, ROIs were manually drawn around the cell of interest, specifically excluding nuclei and autofluorescent gut granules. GA3 was used for threshold-based segmentation of the ER. For experiments in which significant variability in ER fluorescence precluded the use of constant thresholds, thresholds were manually set on blinded images. Images for yeast experiments were deconvolved using softWoRx, and the integrated density of fluorescence intensity units were normalized with cell surface area using Fiji.

### Transmission electron microscopy (TEM) and analysis

Worms were collected from bacterial lawns and washed thrice with M9. Worm pellets were resuspended in 0.15 M sucrose prepared in M9 buffer and loaded into 200 µm deep well of a A-type carrier (Leica). The assembly was covered with flat side of B-type carrier (Leica) and vitrified using a high-pressure freezing machine (Leica EM ICE). The frozen specimens were stored in liquid nitrogen until further processing with a modified freeze-substitution (FS) process^65^. Briefly, FS was performed using a cocktail of 0.5% glutaraldehyde and 0.1% tannic acid in acetone. Samples were transferred into an automatic freeze-substitution machine (AFS2, Leica) pre-cooled to -140 °C. Over 3 h, samples were warmed to -90 °C, and then they were incubated at -90 °C for 114 h. Samples were then washed 4× for 30 min each with acetone. The final acetone wash was replaced with 2% osmium tetroxide in acetone. Following this, samples were warmed to -20 °C over 12 h, after which temperature was maintained at -20 °C for an additional 10 h before further warming to 0 °C over 4 h. Samples were then transferred into an ice filled bath and washed 4× for 30 min each with acetone and infiltrated with Spurr’s resin (Electron Microscopy Sciences). During final steps of resin infiltration, individual worms were transferred from carriers into resin molds with a fine needle, and samples were cured at 60 °C for 72 h. Thin sections (70 nm) were cut using a Leica UC7 and a Diatome diamond knife, positioned on grids, and stained with uranyl acetate and Reynold’s lead citrate. These sections were then imaged on a Tecani T12 TEM operating at 100 keV using an AMT CMOS camera. Amira (Thermo Fisher Scientific) was used for image processing and analysis. Briefly, the hypodermal ER and cytosolic area were manually segmented with the freehand masking tool, and ER footprint and perimeter-area ratio were calculated.

### Scanning electron microscopy (SEM) and image analysis

Brain tissue for SEM imaging was prepared as previously described^66^. Briefly, one male mouse from each age group (6 mo and 18 mo) was euthanized and perfused for 30 s with 37 °C Ringer’s solution (0.79% NaCl, 0.038% KCl, 0.02% MgCl_2_·6H_2_O, 0.018% Na_2_HPO_4_, 0.125% NaHCO_3_, 0.03% CaCl_2_·2H_2_O, 0.2% dextrose, and 1000 U heparin) followed by a transcardiac perfusion with fresh, ice-cold 2.5% glutaraldehyde and 2% PFA in 0.15M sodium cacodylate at 10 mL/min for 10 min. The brains were dissected and sliced sagittally with a vibratome to prepare 150 µm-thick sections and postfixed at 4°C overnight. Brain slices were washed in cold 0.15 M sodium cacodylate buffer and then fixed in 2% osmium tetroxide and 1.5% potassium ferrocyanide in 0.15 M sodium cacodylate buffer for 1 h at room temperature (∼23 °C). Samples were washed several times in double distilled water (ddH_2_0), placed in a 0.5% thiocarbohydrazide solution for 30 min, and then washed 5× in ddH_2_0. Next, the slices were placed in a 2% aqueous osmium tetroxide solution for 1 h, thoroughly washed in ddH_2_0, and transferred to a 2% aqueous uranyl acetate solution and placed at 4 °C overnight. The next day, slices were washed with ddH_2_0 and placed into Walton’s lead aspartate solution for 30 min and baked at 60 °C using a bench-top oven. Baked samples were thoroughly washed with ddH_2_0 followed by a serial dehydration series on ice using ice-cold 70% EtOH, 90% EtOH, 100% EtOH, 100% EtOH, and dry acetone (10 min/step). The brain slices were then subsequently washed in 1:3, 1:1, and 3:1 solutions of Durcupan ACM:acetone for 12 h in each solution. Finally, the slices were incubated in three, 24-hour washes of 100% Durcupan ACM before being baked for 48 hours at 65°C for solidification.

Slices containing layer 2 motor cortical neurons were imaged on a Zeiss Gemini class scanning electron microscope. Images were blinded, and one neural soma from the central region of each image was selected for analysis. The ER and somatic cytosolic area were manually segmented in NIS-AR, and ER footprint was calculated.

### Cloning and transgenesis – C. elegans

To generate extrachromosomal array *wbmEx222[eft-3p::GFP::sec-61.b::unc-54 UTR]*, a *C. elegans* codon-optimized^67^ GFP(S65C,Q80R) sequence was fused to the native *sec-61.b* sequence in an expression vector to generate plasmid pKB22. The plasmid was column purified and microinjected as previously described^68^.

To generate *ret-1::GFP* and *ret-1::mKate2*, codon-optimized GFP(S65C,Q80R) and mKate2 sequences were amplified with an N-terminal GGGGS linker from plasmids pKB51(GFP) and pKB52(mKate2) using common forward primer Fwd (5’-3’): GTG CTC CAG TCG CCG CTG AAG AGA AGA AGG ATC AAG GTG GCG GAG GTT CTG G and specific reverse primers Rev-GFP (5’-3’): caa tgg caa agt gtg ttc ttc ttt caa tcg atT TAT TTG TAT AGT TCA TCC ATG or Rev-mKate2 (5’-3’): caa tgg caa agt gtg ttc ttc ttt caa tcg atT TAA CGG TGT CCG AGC TTG GAT G. The repair templates were column purified and injected as previously described^69^ with a guide (Dharmacon) against the following target sequence (5’-3’): AAT GGC AAA GTG TGT TCG GA. For *ret-1::wrmScarlet11,* an ssODN repair template including the *wrmScarlet11* sequence (5’-3’: GTG CTC CAG TCG CCG CTG AAG AGA AGA AGG ATC AAG GTG GCG GAG GTT CTT ACA CCG TCG TCG AGC AAT ACG AGA AGT CCG TCG CCC GTC ACT GCA CCG GAG GAA TGG ATG AGT TAT ACA AGT AAA TCG ATTGAA AGA AGA ACA CAC TTT GCC ATT GTT TTT C) was synthesized (IDT). The repair template was injected with the same guide used for the GFP and mKate2 fusions. All *ret-1* fusion strains were sequence verified and outcrossed with N2 worms at least four times.

For neuron-specific RET-1 fluorescence, we first generated extrachromosomal array *bugEx10[rab-3p::wrmSct1-10::unc-54 UTR + rab-3p::GFP::unc-54 UTR]* through the cloning and subsequent co-injection of pBK01(∼62 ng/µL) and pBK02(∼53 ng/µL) into N2 worms. To generate plasmid pBK01(*rab-3p::GFP::unc-54 UTR*), the *atf-6* sequence was excised from plasmid pKB41 via BamHI/AgeI digest, and the backbone was re-ligated after Klenow treatment. To generate plasmid pBK02(*rab-3p::wrmSct1-10::unc-54 UTR*), the *wrmSct1-10::unc-54 UTR* sequence was amplified from plasmid pJG100^70^ with primers Fwd (5’-3’): CCC GGG ATG GTA TCG AAG GGA GAA GC and Rev (5’-3’): ACT AGT CTT CCA CTG AGC CTC AAA. The PCR product was ligated into a vector containing the *rab-3p* sequence following XmaI/SpeI digestion. After microinjection, GFP+ worms were crossed with *ret-1::wrmScarlet11* worms to generate strain BUZ112(*ret-1::wrmScarlet11; bugEx10*).

### Cloning and transgenesis – S. cerevisiae

Strains used in this study were derived from SEY6210 cells by fusing fluorescent proteins to the 3’-end of endogenous genes via homologous recombination.

### Protein isolation and SDS-PAGE

Synchronized populations of at least 400 worms were raised as described above on 10 cm NGM plates seeded with 500 µL OP50-1 overnight cultures. To assist with aging a large population, L4 worms expressing 3×FLAG::SEC-61.B were transferred to lawns spotted with 250 µL 5-fluoro-2′-deoxyuridine (FUDR, 1 mg/mL solution) 24 h prior to use. For RET-1::GFP blotting, germline tumors were observed with extremely high levels of RET-1, necessitating a germlineless *glp-1* background. *Ret-1::gfp; glp-1* worms were heatshocked as described above. Worms were collected from bacterial lawns and washed thrice with M9, and worm pellets were flash frozen with liquid nitrogen and stored at -80 °C. Worms were lysed at 4 °C by sonication in RIPA buffer (50 mM Tris, 150 mM NaCl, 1 mM EDTA, 1% Triton X-100, 0.1% sodium deoxycholate, 0.1% SDS, pH 7.5) with protease inhibitors (Roche Applied Science). Lysates were centrifuged at 14,000 g for 15 min at 4 °C, and the supernatants were collected. Protein concentrations were determined via Pierce BCA assay (Thermo Scientific). An equal mass of each sample was diluted in Laemmli Sample Buffer (BioRad) with 2-Mercaptoethanol, heated at 95 °C for 15 min, and resolved via SDS-PAGE electrophoresis.

### Western blotting, protein quantification, and data analysis

Proteins were transferred to activated PVDF membranes over 2 h at 4 °C. Ponceau (Sigma Aldrich) staining, imaging, and washing was conducted per manufacturer recommendations. Target proteins were probed through the following steps: an overnight, 4 °C incubation in blocking solution (5% non-fat milk in TBST); a 2-hour, room temperature incubation in primary antibody solution (5% milk in TBST + antibody (anti-FLAG: 1:1,500 or anti-GFP: 1:1,000)); three, 10-minute, room temperature washes in TBST; and a 1-hour, room temperature incubation in secondary antibody solution (5% milk in TBST + antibody (HRP-conjugated, anti-mouse (1:10,000))). After three more 10-minute, room temperature washes in TBST, membranes were incubated with HRP substrate (Pierce ECL Western Blotting Substrate (Thermo Scientific)) per manufacturer recommendations. All wash and incubation steps were performed with gentle nutation. Ponceau and ECL imaging was performed with an Amersham Imager 600 (GE Life Sciences). Bands were manually selected and quantified in Fiji, and HRP signal was normalized to total protein for analysis.

### Quantification and statistical analysis

Data from published Wcs datasets^35^ were reanalyzed after identifying proteins annotated for ER localization (Wormbase WS292, GO_0005783) and WormCat2.0 annotated functions in either proteostasis (categories: chaperone, protein modification, and stress response) or lipid metabolism. Imaging experiments were conducted in three independent repeats, and data were pooled prior to analysis. For imaging experiments, the experimenter was blinded to experimental conditions prior to either image acquisition or analysis as feasible. Blinding was performed with Fiji script called blind-files, provided as part of the Lab-utility-plugins update site^71^. Isolated comparisons between two, independent groups were conducted via two-tailed, Student’s t-tests. Western blot data were analyzed via two-tailed, paired, ratio t-tests. Categorical morphologic analyses were conducted with Χ^2^ tests. For experiments requiring multiple comparisons, one- or two-way ANOVAs were conducted as appropriate, with post-hoc tests to determine intergroup differences. Prism (v9 & v10, GraphPad) was used for statistical analysis and plot generation. Graph midlines indicate mean ± 95 % CI unless otherwise specified. Statistical tests, p-values, and sample sizes are indicated in figure legends.

## Supplemental data

### Supplemental figures

**Fig. S1.**
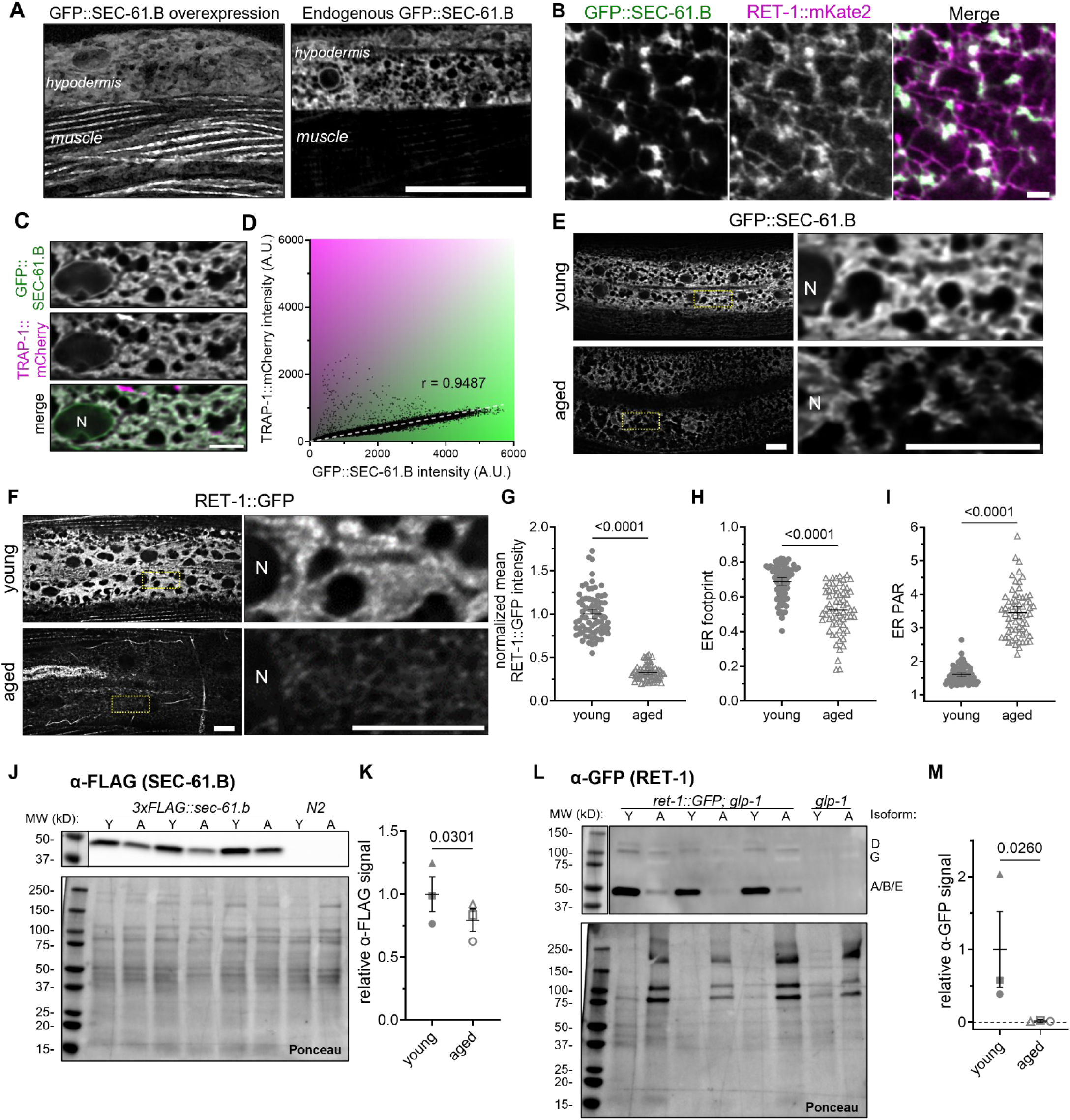
ER subdomain structure and protein expression are reduced in aging, related to Figure 1. (A) Representative fluorescence images of GFP::SEC-61.B overexpression (left) vs. endogenous tagging (right) revealing mislocalization of overexpressed SEC-61.B to muscle SR. Scale: 10 µm. (B) Super-resolution image from a cortical region of the *C. elegans* hypodermis demonstrating expected subdomain enrichment of RET-1::mKate2 in ER tubules and sheet edges and GFP::SEC-61.B to patches of ER sheets. Scale: 2 µm. (C) Representative fluorescence image of hypodermal ER co-labeled with translocon subunits GFP::SEC-61.B and TRAP-1::mCherry,. Scale: 10 µm. (D) Pearson’s correlation (*r*) between GFP::SEC-61.B and TRAP-1::mCherry. r = 0.9487. (E) Hypodermal GFP::SEC-61.Β in young and aged worms. Scale: 10 µm. (F) Hypodermal RET-1::GFP in young and aged worms. Scale: 10 µm. (G-I) Quantification of normalized mean intensity (G), footprint (H), and PAR (I) in young vs. aged RET-1::GFP worms. N = 76 young, 65 aged, pooled across 3 replicates. Analyzed via t-tests. (J) Western blotting (α-FLAG) on lysates from *3xFLAG::sec-61.β* worms, with N2 controls. Ladder annotated with molecular weights. N = 3 young samples (“Y,” d1), 3 aged samples (“A,” d7). Ponceau staining included for total protein normalization. (K) Quantification of α-FLAG signal in young and aged worms, relative to total protein levels (Ponceau) and normalized to d1. Replicates indicated by similar shapes. Analyzed via ratio paired t-test. (L) Western blotting (α-GFP) on lysates from *ret-1::GFP; glp-1* worms, with *glp-1* controls. Ladder annotated with molecular weights. Lower molecular weight bands are consistent with predictions for RET-1.A/B/E; higher molecular weight bands are consistent with predictions for RET-1.D/G. N = 3 young samples (“Y,” d1), 3 aged samples (“A,” d7)). Ponceau staining included for total protein normalization.

**Fig. S2.**
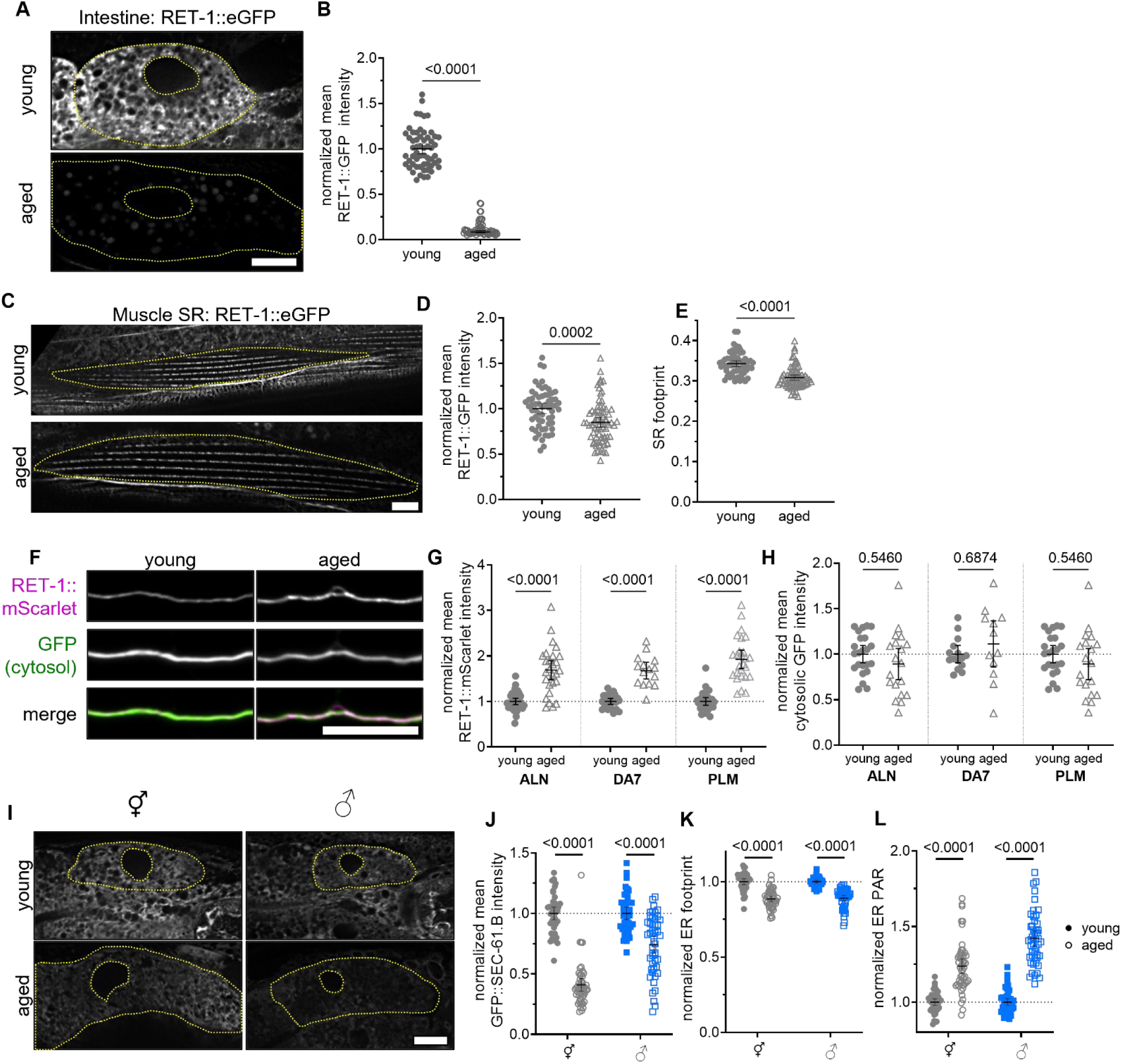
Additional tissues and ER structures undergo age-onset ER remodeling, related to Figure 2. (A) Imaging of intestinal RET-1::GFP in young and aged worms. Scale: 10 µm. (B) Quantification of intestinal RET-1::GFP intensity, normalized to d1. N = 58 young, 59 aged, pooled across 3 replicates. Analyzed via t-test. (C) Imaging of RET-1::GFP of muscle myofilament-associated SR in young and aged animals. Scale: 10 µm. (D-E) Quantification of normalized mean intensity (D), and footprint (E) of RET-1::GFP in muscle SR. N = 61 young, 72 aged, pooled across 3 replicates. Analyzed via t-tests. (F) Maximum intensity projections of neuron-specific RET-1::wrmScarlet in the neurites of the ALN neuron marked with cytoplasmic GFP. Scale = 10 µm. (G-H) Quantification of mean RET-1::wrmScarlet (G) and cytoplasmic GFP (H) intensity along the ALN, DA7, and PLM neurites in young and aged worms. For (G) N_ALN_ = 34 young, 27 aged; N_DA7_ = 23 young, 15 aged; N_PLM_ = 28 young, 26 aged; pooled across 3 replicates. For (H), N_ALN_ = 23 young, 19 aged; N_DA7_ = 15 young, 12 aged; N_PLM_ = 23 young, 19 aged; pooled across 2 replicates. Analyzed via one-way ANOVAs with post hoc Šidák tests. (I) Imaging of intestinal GFP::SEC-61.Β during aging of hermaphrodite (⚥) and male (♂) worms. Scale: 10 µm. (J-L) Quantification of mean intensity (J), footprint (K), and PAR (L) in intestine of GFP::SEC-61.B worms across age, normalized to d1 for each sex. N_⚥_ = 42 young, 50 aged; N_♂_ = 43 young, 49 aged; pooled across 3 replicates. Analyzed via t-tests. For all graphs, error bars indicate mean ± 95% CI. For images, “N” = nucleus.

**Fig. S3.**
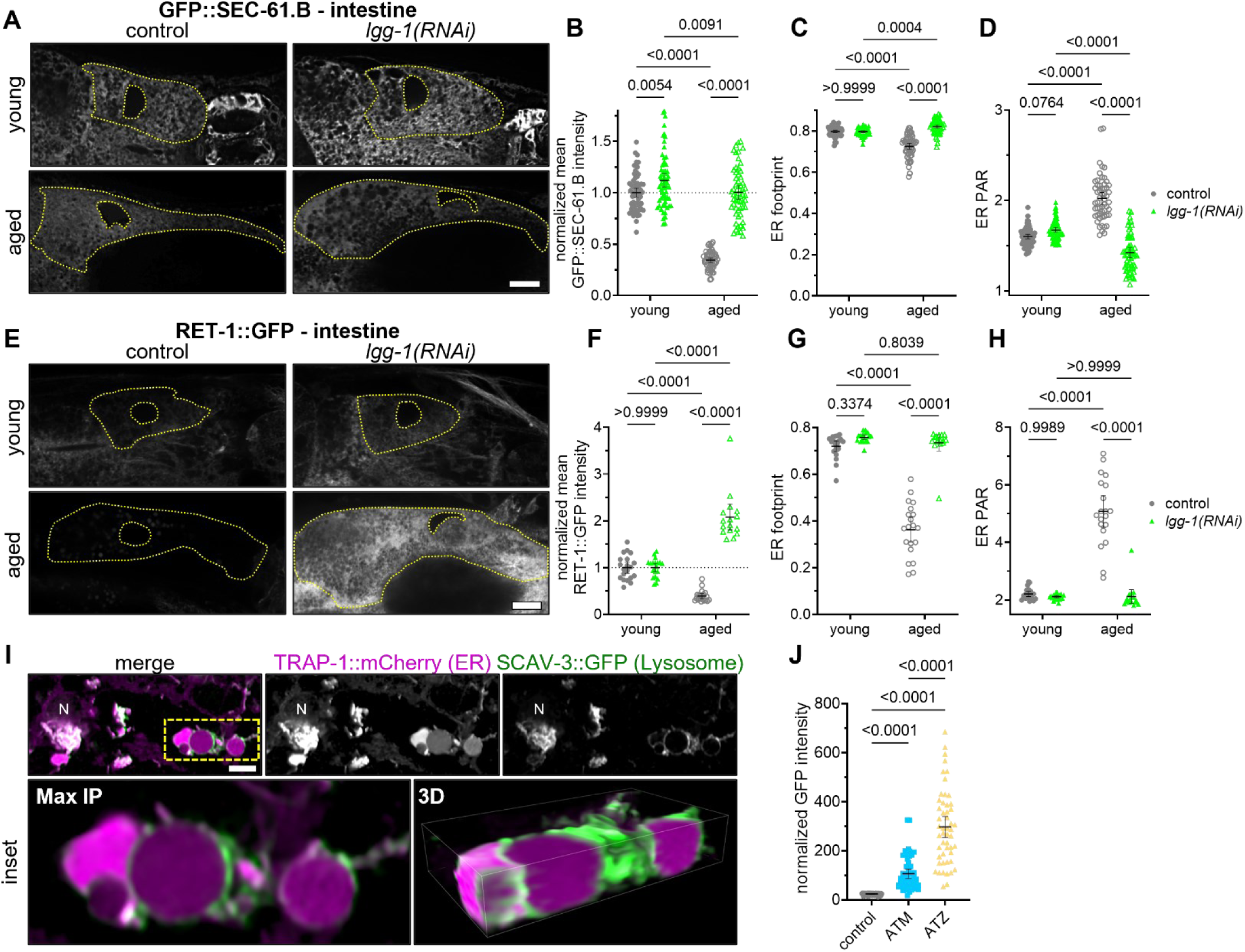
Autophagy regulates the turnover of multiple ER subdomain proteins, related to Figure 3. (A) Intestinal GFP::SEC-61.Β in young and aged control (e.v.) and *lgg-1(RNAi)* worms. All scales: 10 µm. (B-D) Intestinal GFP::SEC-61.B normalized mean intensity (B), footprint (C), and PAR (D) in young and aged control and *lgg-1* KD worms. Young = filled shapes; aged = hollow shapes. N_e.v._ = 63 young, 55 aged; N*_lgg-1_* = 64 young, 58 aged, pooled across 3 replicates. Analyzed via two-way ANOVA and post hoc Šidák tests. (E) Intestinal RET-1::GFP in young and aged control, *unc-51* KD, and *lgg-1* KD worms. All scales: 10 µm. (F-H) Intestinal RET-1::GFP normalized mean intensity (F), footprint (G), and PAR (H) in young and aged control and *lgg-1* KD worms. Young = filled shapes; aged = hollow shapes. N_e.v._ = 20 young, 20 aged; N*_lgg-1_* = 20 young, 16 aged; from 1 replicate. Analyzed via two-way ANOVA and post hoc Šidák tests. (I) Representative fluorescence image shows accumulation of ER reporter TRAP-1::mCherry within lysosomes (SCAV-3::GFP) during early aging. Inset displayed as maximum intensity and 3-D projections. Scale: 5 µm. (M) Intestinal GFP::AT intensity in control, GFP::ATM-, and GFP::ATZ-expressing worms. N = 51 worms/group, pooled across 3 replicates. One way ANOVA, p < 0.0001, post hoc Tukey test. For all graphs, error bars indicate mean ± 95% CI. For images, “N” = nucleus.

**Fig. S4.**
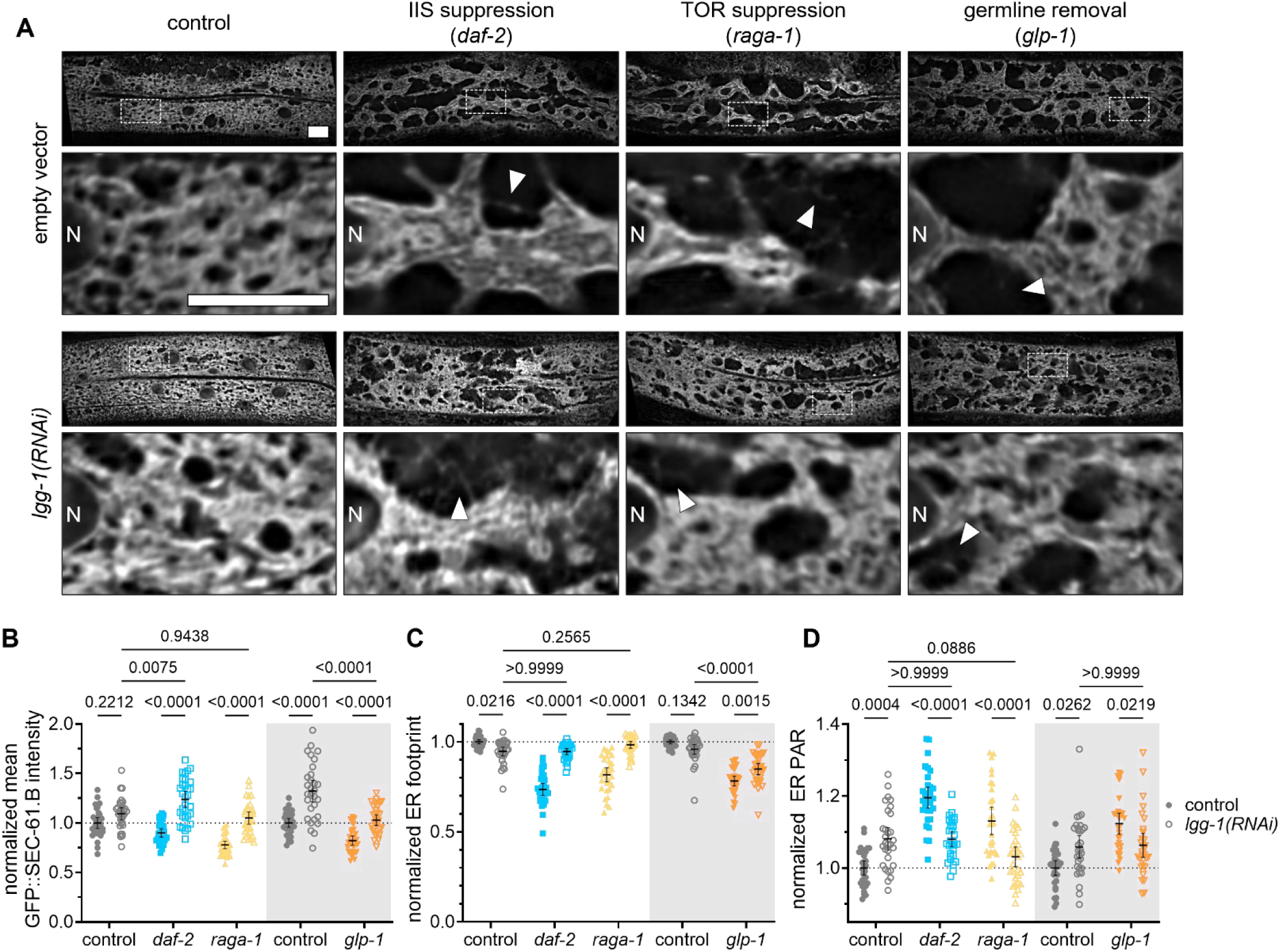
ER remodeling in lifespan-extending conditions depends on autophagy, related to Figure 4. (A) Confocal imaging of hypodermal GFP::SEC-61.Β in young worms from wild type, *daf-*2, *raga-1*, and *glp-1* backgrounds fed control or *lgg-1* dsRNA. All scales: 10 µm. “N” = nucleus. Arrows indicate sparse ER tubules. (B-D) Quantification of mean intensity (C), footprint (D), and PAR (E) of GFP::SEC-61.B in long-lived animals relative to wild type controls after feeding of empty vector or *lgg-1* dsRNA. Colored backgrounds group experimental conditions with their respective controls. N = 30 control, 30 *daf-2*, 30 *raga-1*, 30 control_(25°C)_, 30 *glp-1*, pooled across 3 replicates. Two-way ANOVA, p < 0.0001, post hoc Šidák test.

### Supplemental tables

**Supplemental Table 1:** yeast chronological lifespan data.

| strain | autophagy? | ER-phagy? | replicate | median lifespan (d) | 95% CI |
| --- | --- | --- | --- | --- | --- |
| control | + | + | 1 | 12.2 | 7.0-22.4 |
|  |  |  | 2 | 13.0 | 8.0-22.2 |
|  |  |  | 3 | 12.4 | 7.4-21.8 |
|  |  |  | 4 | 12.3 | 8.3-18.8 |
|  |  |  | Average | 12.5 |  |
| *Δatg5* | - | + | 1 | 5.6 | 3.1-9.4 |
|  |  |  | 2 | 5.4 | 3.3-8.3 |
|  |  |  | 3 | 5.1 | 3.3-7.4 |
|  |  |  | 4 | 5.6 | 3.1-9.4 |
|  |  |  | Average | 5.4 |  |
| *Δatg39* | + | - | 1 | 6.6 | 3.7-11.0 |
|  |  |  | 2 | 5.8 | 3.5-8.9 |
|  |  |  | 3 | 6.3 | 4.0-9.4 |
|  |  |  | 4 | 6.2 | 4.2-8.8 |
|  |  |  | Average | 6.2 |  |
| *Δatg40* | + | - | 1 | 6.6 | 4.0-10.3 |
|  |  |  | 2 | 6.1 | 4.3-8.5 |
|  |  |  | 3 | 6.4 | 4.0-9.7 |
|  |  |  | 4 | 6.1 | 4.0-8.8 |
|  |  |  | Average | 6.3 |  |
| *Δsec66* | + | - | 1 | 6.4 | 4.8-8.5 |
|  |  |  | 2 | 6.4 | 4.4-9.1 |
|  |  |  | 3 | 6.1 | 3.9-9.1 |
|  |  |  | 4 | 6.5 | 4.2-9.8 |
|  |  |  | Average | 6.4 |  |
| Šidák multiple comparison test: |  |  |  |  |  |
| Comparison |  | *p*<sub>adj</sub> | Comparison |  | *p*<sub>adj</sub> |
| control vs. *Δatg5* |  | <0.0001 | *Δatg5* vs. *Δatg40* |  | 0.0066 |
| control vs. *Δatg39* |  | <0.0001 | *Δatg5* vs. *Δsec66* |  | 0.0035 |
| control vs. *Δatg40* |  | <0.0001 | *Δatg39* vs. *Δatg40* |  | >0.9999 |
| control vs. *Δsec66* |  | <0.0001 | *Δatg39* vs. *Δsec66* |  | 0.9995 |
| *Δatg5* vs. *Δatg39* |  | 0.0127 | *Δatg40* vs. *Δsec66* |  | >0.9999 |

**Supplemental Table 2:**
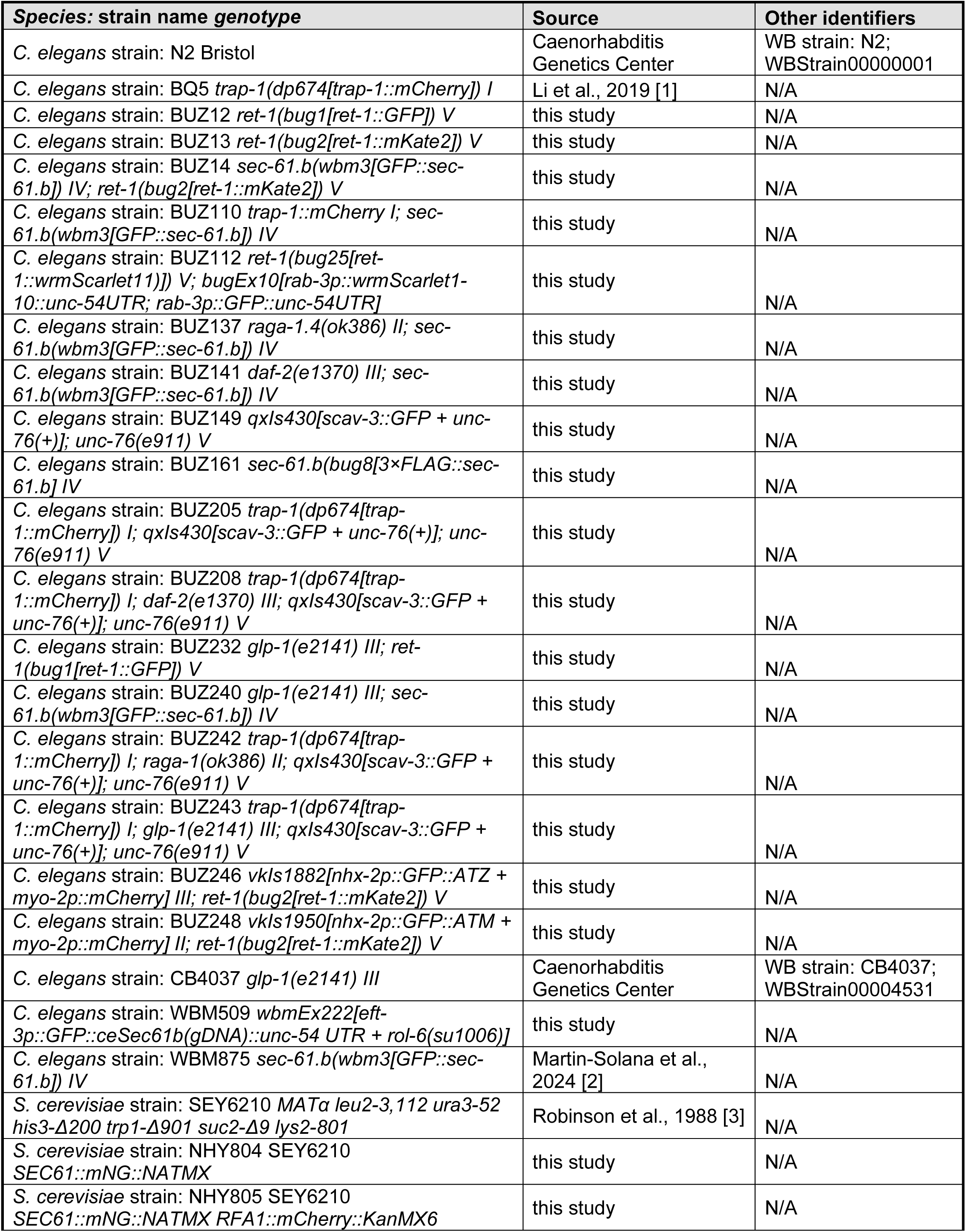
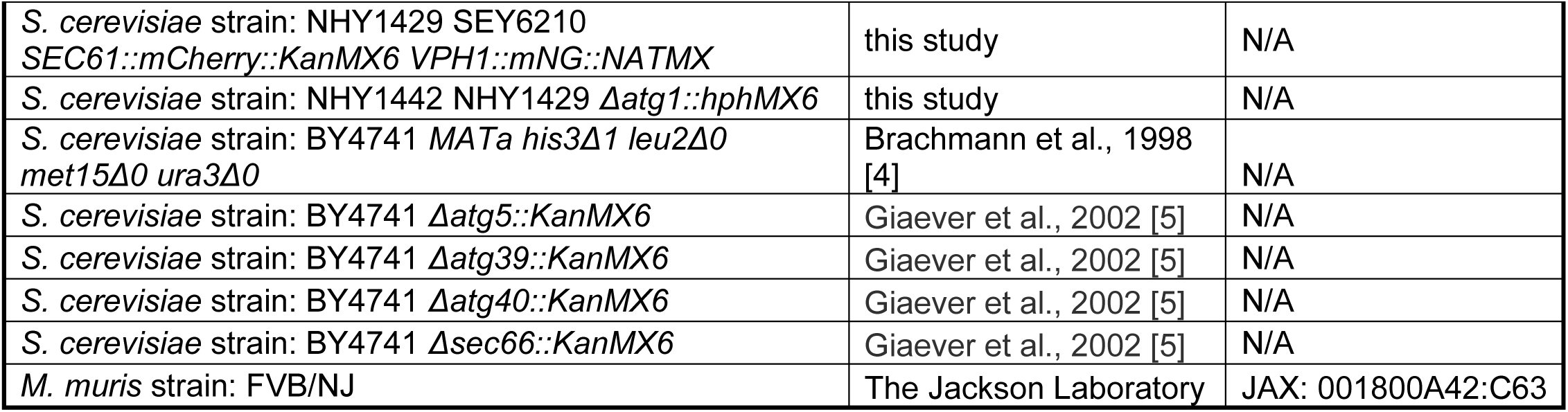
model organisms used in this study.

---

**Supplemental Table 3:**
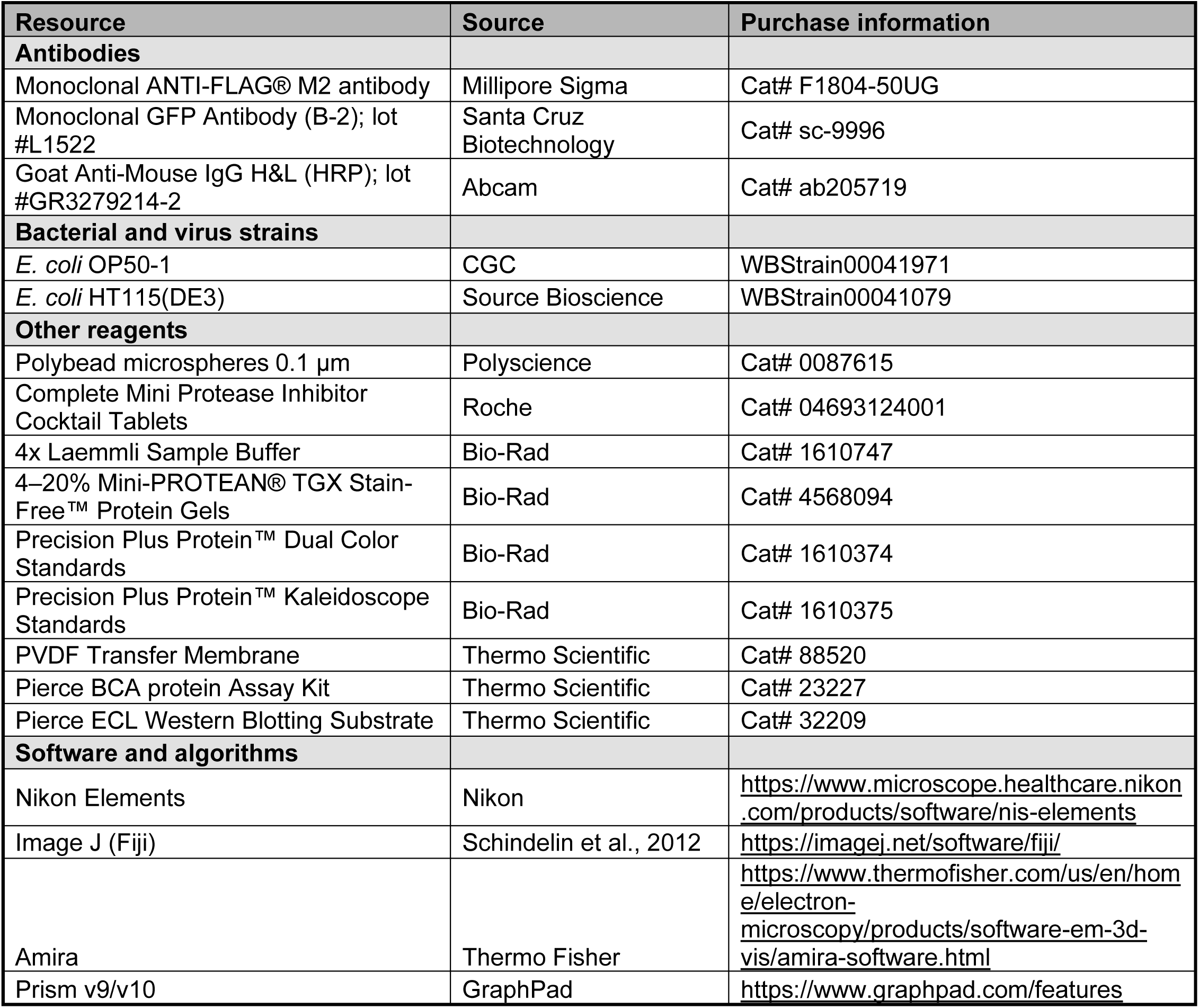
key reagents used.

## References

1. Metcalf, M. G., Higuchi-Sanabria, R., Garcia, G., Tsui, C. K. & Dillin, A. Beyond the cell factory: Homeostatic regulation of and by the UPRER. Sci Adv 6, eabb9614 (2020).

2. Arruda, A. P. & Parlakgül, G. Endoplasmic Reticulum Architecture and Inter-Organelle Communication in Metabolic Health and Disease. Cold Spring Harb. Perspect. Biol. (2022) doi:10.1101/cshperspect.a041261.

3. Phillips, M. J. & Voeltz, G. K. Structure and function of ER membrane contact sites with other organelles. Nat. Rev. Mol. Cell Biol. 17, 69–82 (2016).

4. Westrate, L. M., Lee, J. E., Prinz, W. A. & Voeltz, G. K. Form follows function: the importance of endoplasmic reticulum shape. Annu. Rev. Biochem. 84, 791–811 (2015).

5. Hotamisligil, G. S. Endoplasmic reticulum stress and the inflammatory basis of metabolic disease. Cell 140, 900–917 (2010).

6. Mandl, J., Mészáros, T., Bánhegyi, G., Hunyady, L. & Csala, M. Endoplasmic reticulum: nutrient sensor in physiology and pathology. Trends Endocrinol. Metab. 20, 194–201 (2009).

7. López-Otín, C., Blasco, M. A., Partridge, L., Serrano, M. & Kroemer, G. Hallmarks of aging: An expanding universe. Cell 186, 243–278 (2023).

8. Donahue, E. K. F., Ruark, E. M. & Burkewitz, K. Fundamental roles for inter-organelle communication in aging. Biochem. Soc. Trans. 50, 1389–1402 (2022).

9. Walter & Ron. REVIEWS The Unfolded Protein Response: From Stress Pathway to Homeostatic Regulation. Science 334, (2011).

10. De-Souza, E. A., Cummins, N. & Taylor, R. C. IRE-1 endoribonuclease activity declines early in C. elegans adulthood and is not rescued by reduced reproduction. Front Aging 3, 1044556 (2022).

11. Rabek, J. P., Boylston, W. H., 3rd & Papaconstantinou, J. Carbonylation of ER chaperone proteins in aged mouse liver. Biochem. Biophys. Res. Commun. 305, 566–572 (2003).

12. Nuss, J. E., Choksi, K. B., DeFord, J. H. & Papaconstantinou, J. Decreased enzyme activities of chaperones PDI and BiP in aged mouse livers. Biochem. Biophys. Res. Commun. 365, 355–361 (2008).

13. Erickson, R. R., Dunning, L. M. & Holtzman, J. L. The effect of aging on the chaperone concentrations in the hepatic, endoplasmic reticulum of male rats: the possible role of protein misfolding due to the loss of chaperones in the decline in physiological function seen with age. J. Gerontol. A Biol. Sci. Med. Sci. 61, 435–443 (2006).

14. Hotamisligil, G. S. Inflammation, metaflammation and immunometabolic disorders. Nature 542, 177–185 (2017).

15. Franceschi, C., Garagnani, P., Parini, P., Giuliani, C. & Santoro, A. Inflammaging: a new immune-metabolic viewpoint for age-related diseases. Nat. Rev. Endocrinol. 14, 576–590 (2018).

16. Wang, X., Li, S., Wang, H., Shui, W. & Hu, J. Quantitative proteomics reveal proteins enriched in tubular endoplasmic reticulum of Saccharomyces cerevisiae. Elife 6, (2017).

17. Lak, B. et al. Specific subdomain localization of ER resident proteins and membrane contact sites resolved by electron microscopy. Eur. J. Cell Biol. 100, 151180 (2021).

18. Zheng, P. et al. DNA damage triggers tubular endoplasmic reticulum extension to promote apoptosis by facilitating ER-mitochondria signaling. Cell Res. 28, 833–854 (2018).

19. Zheng, P. et al. ER proteins decipher the tubulin code to regulate organelle distribution. Nature 601, 132–138 (2022).

20. Parlakgül, G. et al. Regulation of liver subcellular architecture controls metabolic homeostasis. Nature 603, 736–742 (2022).

21. Blackstone, C., O’Kane, C. J. & Reid, E. Hereditary spastic paraplegias: membrane traffic and the motor pathway. Nat. Rev. Neurosci. 12, 31–42 (2011).

22. Zhu, P.-P. et al. Transverse endoplasmic reticulum expansion in hereditary spastic paraplegia corticospinal axons. Hum. Mol. Genet. 31, 2779–2795 (2022).

23. Sree, S., Parkkinen, I., Their, A., Airavaara, M. & Jokitalo, E. Morphological Heterogeneity of the Endoplasmic Reticulum within Neurons and Its Implications in Neurodegeneration. Cells 10, (2021).

24. Shibata, Y. et al. Mechanisms determining the morphology of the peripheral ER. Cell 143, 774–788 (2010).

25. Voeltz, G. K., Prinz, W. A., Shibata, Y., Rist, J. M. & Rapoport, T. A. A class of membrane proteins shaping the tubular endoplasmic reticulum. Cell 124, 573–586 (2006).

26. Puhka, M., Vihinen, H., Joensuu, M. & Jokitalo, E. Endoplasmic reticulum remains continuous and undergoes sheet-to-tubule transformation during cell division in mammalian cells. J. Cell Biol. 179, 895–909 (2007).

27. Ferro-Novick, S., Reggiori, F. & Brodsky, J. L. ER-Phagy, ER Homeostasis, and ER Quality Control: Implications for Disease. Trends Biochem. Sci. 46, 630–639 (2021).

28. Chen, S. et al. Vps13 is required for the packaging of the ER into autophagosomes during ER-phagy. Proc. Natl. Acad. Sci. U. S. A. 117, 18530–18539 (2020).

29. Liang, J. R. et al. A Genome-wide ER-phagy Screen Highlights Key Roles of Mitochondrial Metabolism and ER-Resident UFMylation. Cell 180, 1160–1177.e20 (2020).

30. Rolls, M. M., Hall, D. H., Victor, M., Stelzer, E. H. K. & Rapoport, T. A. Targeting of rough endoplasmic reticulum membrane proteins and ribosomes in invertebrate neurons. Mol. Biol. Cell 13, 1778–1791 (2002).

31. Shibata, Y. et al. The reticulon and DP1/Yop1p proteins form immobile oligomers in the tubular endoplasmic reticulum. J. Biol. Chem. 283, 18892–18904 (2008).

32. Li, X. et al. Requirement for translocon-associated protein (TRAP) α in insulin biogenesis. Sci Adv 5, eaax0292 (2019).

33. Haithcock, E. et al. Age-related changes of nuclear architecture in Caenorhabditis elegans. Proc. Natl. Acad. Sci. U. S. A. 102, 16690–16695 (2005).

34. Scaffidi, P. & Misteli, T. Lamin A-dependent nuclear defects in human aging. Science 312, 1059–1063 (2006).

35. Walther, D. M. et al. Widespread Proteome Remodeling and Aggregation in Aging C. elegans. Cell 161, 919–932 (2015).

36. Cuentas-Condori, A. et al. C. elegans neurons have functional dendritic spines. Elife 8, (2019).

37. Ezcurra, M. et al. C. elegans Eats Its Own Intestine to Make Yolk Leading to Multiple Senescent Pathologies. Curr. Biol. 28, 2544–2556.e5 (2018).

38. Fregno, I. & Molinari, M. Proteasomal and lysosomal clearance of faulty secretory proteins: ER-associated degradation (ERAD) and ER-to-lysosome-associated degradation (ERLAD) pathways. Crit. Rev. Biochem. Mol. Biol. 54, 153–163 (2019).

39. Hayashi, Y. et al. TOLLIP acts as a cargo adaptor to promote lysosomal degradation of aberrant ER membrane proteins. EMBO J. 42, e114272 (2023).

40. Costantini, L. M. et al. A palette of fluorescent proteins optimized for diverse cellular environments. Nat. Commun. 6, 7670 (2015).

41. Hipp, M. S., Kasturi, P. & Hartl, F. U. The proteostasis network and its decline in ageing. Nat. Rev. Mol. Cell Biol. 20, 421–435 (2019).

42. Teckman, J. H. & Perlmutter, D. H. Retention of mutant α1-antitrypsin Z in endoplasmic reticulum is associated with an autophagic response. American Journal of Physiology-Gastrointestinal and Liver Physiology 279, G961–G974 (2000).

43. Schuck, S., Gallagher, C. M. & Walter, P. ER-phagy mediates selective degradation of endoplasmic reticulum independently of the core autophagy machinery. J. Cell Sci. 127, 4078–4088 (2014).

44. Beese, C. J., Brynjólfsdóttir, S. H. & Frankel, L. B. Selective Autophagy of the Protein Homeostasis Machinery: Ribophagy, Proteaphagy and ER-Phagy. Frontiers in Cell and Developmental Biology 7, (2020).

45. Foronda, H. et al. Heteromeric clusters of ubiquitinated ER-shaping proteins drive ER-phagy. Nature (2023) doi:10.1038/s41586-023-06090-9.

46. González, A. et al. Ubiquitination regulates ER-phagy and remodelling of endoplasmic reticulum. Nature (2023) doi:10.1038/s41586-023-06089-2.

47. Long, O. S. et al. A C. elegans model of human α1-antitrypsin deficiency links components of the RNAi pathway to misfolded protein turnover. Hum. Mol. Genet. 23, 5109–5122 (2014).

48. Mochida, K. et al. Receptor-mediated selective autophagy degrades the endoplasmic reticulum and the nucleus. Nature 522, 359–362 (2015).

49. Walther, D. M. et al. Widespread Proteome Remodeling and Aggregation in Aging C. elegans. Cell 168, 944 (2017).

50. Ge, L. et al. Remodeling of ER-exit sites initiates a membrane supply pathway for autophagosome biogenesis. EMBO Rep. 18, 1586–1603 (2017).

51. Tan, J. X. & Finkel, T. A phosphoinositide signalling pathway mediates rapid lysosomal repair. Nature 609, 815–821 (2022).

52. Austad, S. N. & Hoffman, J. M. Is antagonistic pleiotropy ubiquitous in aging biology? Evol Med Public Health 2018, 287–294 (2018).

53. Gladyshev, V. N. Aging: progressive decline in fitness due to the rising deleteriome adjusted by genetic, environmental, and stochastic processes. Aging Cell 15, 594–602 (2016).

54. Villalobos, T. V. et al. Tubular lysosome induction couples animal starvation to healthy aging. Nat Aging 3, 1091–1106 (2023).

55. Sharma, A., Smith, H. J., Yao, P. & Mair, W. B. Causal roles of mitochondrial dynamics in longevity and healthy aging. EMBO Rep. 20, e48395 (2019).

56. Burkewitz, K. et al. Atf-6 Regulates Lifespan through ER-Mitochondrial Calcium Homeostasis. Cell Rep. 32, 108125 (2020).

57. Taylor, R. C. & Dillin, A. XBP-1 is a cell-nonautonomous regulator of stress resistance and longevity. Cell 153, 1435–1447 (2013).

58. Daniele, J. R. et al. UPRER promotes lipophagy independent of chaperones to extend life span. Sci Adv 6, eaaz1441 (2020).

59. Nguyen, T. T. & Corvera, S. Adipose tissue as a linchpin of organismal ageing. Nat Metab 6, 793–807 (2024).

60. Palikaras, K. et al. Ectopic fat deposition contributes to age-associated pathology in Caenorhabditis elegans. J. Lipid Res. 58, 72–80 (2017).

61. Diaz-Ruiz, A., Price, N. L., Ferrucci, L. & de Cabo, R. Obesity and lifespan, a complex tango. Sci. Transl. Med. 15, eadh1175 (2023).

62. Stiernagle, T. Maintenance of C. Elegans. (WormBook, 2006).

63. Curran, S. P. & Ruvkun, G. Lifespan regulation by evolutionarily conserved genes essential for viability. PLoS Genet. 3, e56 (2007).

64. Thomas, L. S. V., Schaefer, F. & Gehrig, J. Fiji plugins for qualitative image annotations: routine analysis and application to image classification. F1000Res. **9**, 1248 (2020).

65. Bélanger, S. et al. A versatile enhanced freeze-substitution protocol for volume electron microscopy. Front Cell Dev Biol 10, 933376 (2022).

66. Toyama, B. H. et al. Visualization of long-lived proteins reveals age mosaicism within nuclei of postmitotic cells. J. Cell Biol. 218, 433–444 (2019).

67. Redemann, S. et al. Codon adaptation-based control of protein expression in C. elegans. Nat. Methods 8, 250–252 (2011).

68. Evans, T. C. Transformation and microinjection. in WormBook (ed. The C. elegans Research Community) (2006).

69. Paix, A., Folkmann, A., Rasoloson, D. & Seydoux, G. High Efficiency, Homology-Directed Genome Editing in Caenorhabditis elegans Using CRISPR-Cas9 Ribonucleoprotein Complexes. Genetics 201, 47–54 (2015).

70. Goudeau, J. et al. Split-wrmScarlet and split-sfGFP: tools for faster, easier fluorescent labeling of endogenous proteins in Caenorhabditis elegans. Genetics 217, (2021).

71. George, N. M., Gentile Polese, A., Merle, L., Macklin, W. B. & Restrepo, D. Excitable Axonal Domains Adapt to Sensory Deprivation in the Olfactory System. J. Neurosci. 42, 1491–1509 (2022).

## Supplemental Table 2 References

1. Li, X. et al. Requirement for translocon-associated protein (TRAP) α in insulin biogenesis. Sci Adv 5, eaax0292 (2019).

2. Martin-Solana, E. et al. Ribosome-Associated Vesicles promote activity-dependent local translation. bioRxiv (2024) doi:10.1101/2024.06.07.598007.

3. Robinson, J. S., Klionsky, D. J., Banta, L. M. & Emr, S. D. Protein sorting in Saccharomyces cerevisiae: isolation of mutants defective in the delivery and processing of multiple vacuolar hydrolases. Mol. Cell. Biol. 8, 4936–4948 (1988).

4. Brachmann, C. B. et al. Designer deletion strains derived from Saccharomyces cerevisiae S288C: a useful set of strains and plasmids for PCR-mediated gene disruption and other applications. Yeast 14, 115–132 (1998).

5. Giaever, G. et al. Functional profiling of the Saccharomyces cerevisiae genome. Nature 418, 387–391 (2002).

